# A modular encapsulation system for precision delivery of proteins, nucleic acids and small molecules

**DOI:** 10.1101/2024.08.22.607124

**Authors:** Thai D. Luong, Nick Martel, James Rae, Harriet P. Lo, Ye-Wheen Lim, Yeping Wu, Kerrie-Ann McMahon, Haolan Sun, Nick Fletcher, Kristofer Thurecht, Angus P.R. Johnston, Nicholas Ariotti, Tom E. Hall, Robert G. Parton

## Abstract

Targeted nanoparticles have the potential to revolutionize therapeutics for medical applications. Here, we demonstrate the utility of a flexible precision nanovesicle delivery system for functional delivery of DNA, RNA, proteins and drugs into target cells. Nanovesicles generated by the membrane sculpting protein caveolin, termed caveospheres, can be loaded with RNA, DNA, proteins or drugs post-synthesis or incorporate genetically-encoded cargo proteins during production without the need for protein purification. Functionalized fluorescently-labeled caveospheres form a modular system that shows high stability in biological fluids, specific uptake by target-positive cells, and can deliver proteins, drugs, DNA, and mRNA directly to the cytoplasm and nuclei of only the target cells. The negligible level of off-target transduction and uniform levels of targeted expression demonstrates advantages of the system over lipid-mediated gene delivery. Caveospheres can also be engineered to mimic viral particles by displaying the SARS-CoV-2-RBD protein, enabling targeted delivery to human bronchial epithelial cells. We demonstrate their application as a targeted transfection system for cells in culture, and critically, their efficacy in precision tumor killing *in vivo*.

## MAIN TEXT

Targeted delivery in nanomedicine is crucial for maximizing therapeutic efficacy by minimizing off-target impacts and minimizing potential side-effects. Techniques for nanoparticle synthesis and their subsequent functionalization by ligand conjugation often require complex manufacturing processes. ^1, 2^ In the case of antibody-based targeting, binding to the synthesized nanoparticle can be inefficient due to the presence of multiple functional groups on the antibody resulting in heterogeneous orientations. ^3^ In complex diseases, like cancer, diverse therapeutic molecules targeting multiple aspects of cancer biology are necessary. Relying solely on a single receptor or pathway can lead to the expansion of drug-resistant cancer cells. ^4^ This obviates the need for a platform of flexible and versatile yet specific targeted vehicles that can deliver therapeutic molecules to a variety of different cancer cell types. An ideal system would be one that allows for multiple treatment strategies including chemotherapies, immunotherapies, and RNA therapies.

We have developed a modular nanovesicle system based on expression of the mammalian caveolin-1 protein (hereby termed caveolin) in a bacterial host. Caveolin is incorporated into the cytoplasmic membrane of *E.coli* and where it oligomerizes to induce curvature and drive inward vesicle formation **(Scheme 1; Figure S1a)**. ^5^ The resulting nanovesicles (termed caveospheres), of approximately 50nm diameter **(Figure S1b)**, accumulate to an extremely high density within the cytoplasm of the bacteria and can be purified with high yield. Each caveosphere contains approximately 150 caveolin molecules arranged as disc-like oligomers occupying the cytoplasmic leaflet of the vesicles as shown by cryoelectron microscopy, ^5^ and a precisely defined lipid composition. Bacterial proteins are largely excluded from these nanovesicles. ^6^ This structurally-defined and genetically-encoded system allows functional modification of the caveolin fusion proteins. Moieties with defined roles (e.g. antibody-binding, purification) can be attached to both N- and C-termini of the caveolin protein which are both orientated towards the outside of the caveosphere and so are available for interactions with targets. ^7^

Here, we demonstrate the ability of the caveosphere system to precisely deliver encapsulated cargoes into the cytoplasm and nucleus of targeted cells whereby they can effectively modulate cellular responses through delivery of their molecular payload **(Scheme 1)**. ^5^ Specifically, we show i) efficient loading with diverse macromolecules including RNA, DNA, protein and both water-soluble and lipophilic drugs, ii) loaded, functionalized caveospheres can deliver cargo specifically to a variety of different target-positive cells in mixed culture, iii) caveosphere-mediated targeted transfection of RNA or DNA results in high-efficiency, uniform expression in target cells, iv) functionalized, far-labelled caveospheres loaded with chemotherapeutics can be specifically targeted to tumours and are able to reduce tumour size in a mouse xenograft model.

**Scheme 1.**
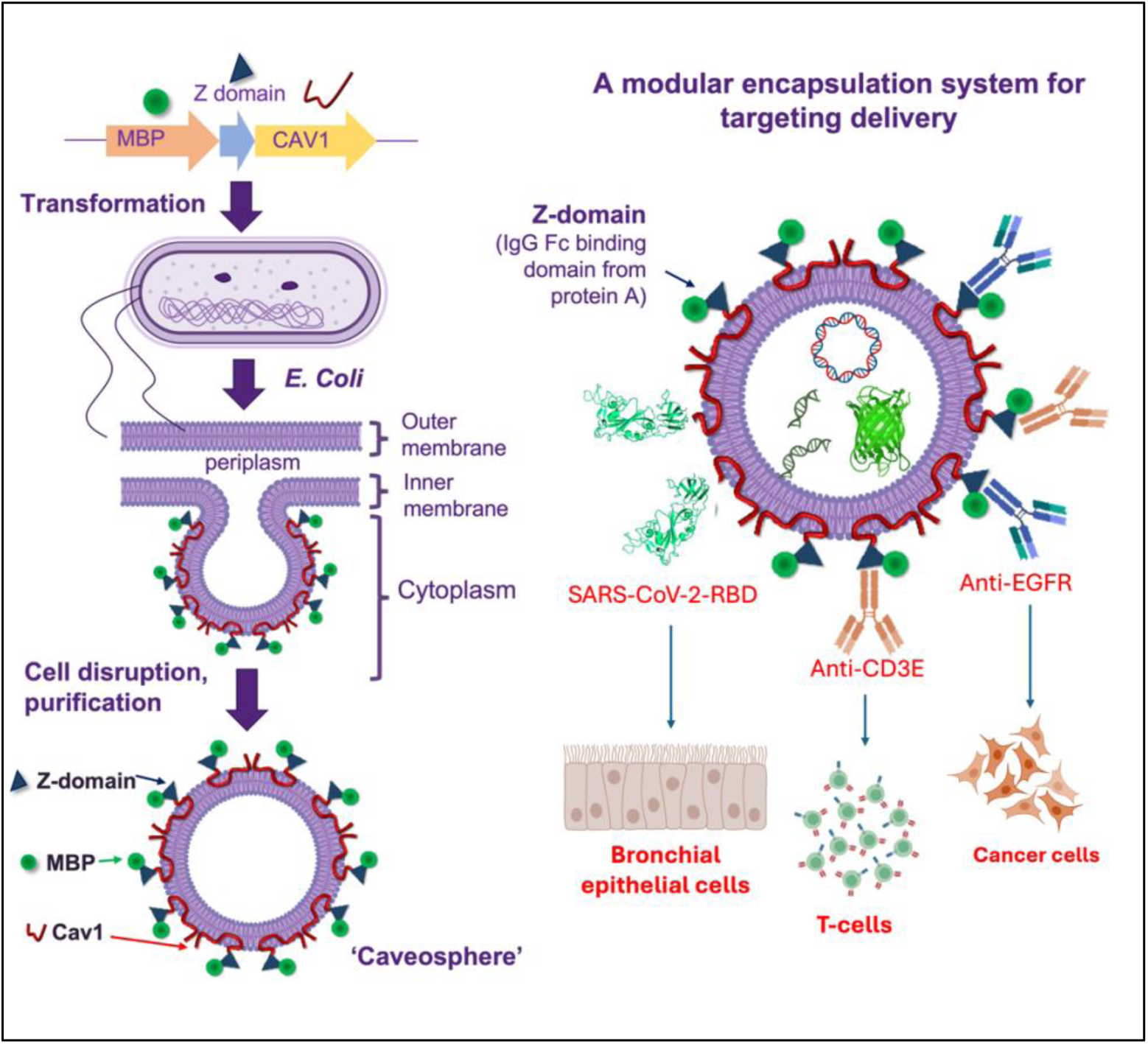
Design and assembly of scalable and tuneable lipid nanoparticles encapsulating diverse cargoes for precise delivery to the cytosol/nuclei of target cells *in vitro* and *in vivo*.

## Results

### Engineering caveospheres for encapsulation of diverse cargoes and cell-specific targeting

We first investigated the possibility of incorporating recombinant proteins into caveospheres as they form by budding into the cytoplasm from the cytoplasmic membrane using a method that avoids cargo protein expression, purification, and encapsulation. We designed a bicistronic vector encoding both the caveosphere-generating fusion protein and the bright fluorescent protein mScarlet (cargo) tagged at the N-terminus with a short periplasmic targeting signal **(Figure 1a)**. The integration of mScarlet into the caveospheres during formation using this genetically-encoded method (termed GE-caveospheres) was demonstrated by their fluorescence post-purification **(Figure 1b i-ii)**. TEM analysis showed GE-caveospheres to possess the typical spherical morphology with a diameter of approximately 50 nm, indistinguishable from unloaded caveospheres generated with the original vector **(Figure 1b iii, Figure S1b)**. Ultracentrifugation of the GE-caveospheres resulted in a brightly fluorescent pellet, which confirmed the association between mScarlet and caveospheres. In contrast, detergent exposure fully solubilized the caveosphere membranes, leading to release of the encapsulated cargo and, consequently, no pellet formation **(Figure S2a-b)**. Encapsulation was further tested by treatment with CuCl_2_ to quench accessible mScarlet fluorescence. Detergent-treated spheres, but not native spheres, showed quenching of the putative cargo protein **(Figure S2c-e)**.

**Figure 1.**
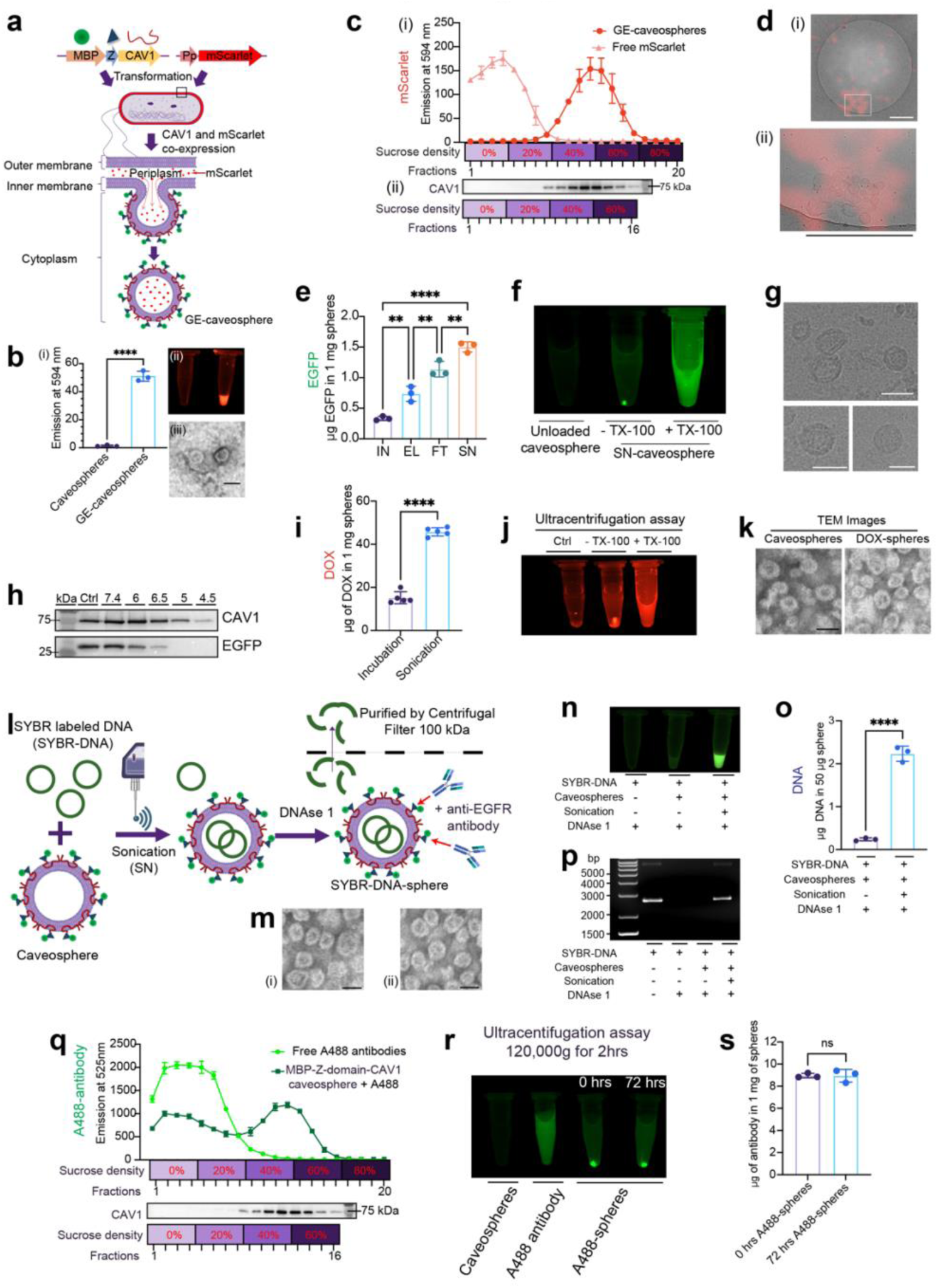
Engineering caveospheres for encapsulation of diverse cargoes and cell-specific targeting. **a**, Development of a genetically-encoded method for loading mScarlet into caveospheres using a periplasmic signal peptide in E. coli, termed “GE-caveospheres”. **b**, (i) Quantification of fluorescence emission at 594 nm of unloaded caveospheres and GE-caveospheres (n=3 technical replicates, Student’s t-test). (ii) Fluorescence emission of unloaded caveospheres (left) and mScarlet-GE-caveospheres (right). (iii) TEM image of GE-caveospheres. Scale bar, 50 nm. **c**, (i) Quantification of fluorescence emission at 594 nm of GE-caveospheres separated by discontinuous sucrose gradient (0%, 20%, 40%, 60%, and 80%, 4 fractions per concentration). Free mScarlet protein was used as a control; n=3 technical replicates. (ii) Western analysis for MBP-Z-domain-CAV1 (72kDa) using anti-CAV1 antibody in fractions collected after a discontinuous sucrose gradient assay of GE-caveospheres (0%, 20%, 40%, and 60%, 4 fractions per concentration). Molecular weight marker (75kDa) as indicated. **d**, (i) Cryo-TEM of mScarlet-GE-caveosphere. (ii) Magnification of boxed area in (i). Scale bars, 500 nm. **e**, Comparison of three physical methods (sonication [SN], electroporation [EL] and freeze-thaw cycling [FT]) for encapsulation of EGFP into caveospheres using fluorescence emission of EGFP-loaded caveospheres (at 507 nm shown as µg protein/mg spheres; n=3 technical replicates, one-way ANOVA, Tukey’s multiple comparison test). IN=incubation of proteins and caveospheres without physical treatment. **f**, Fluorescence emission of EGFP-loaded caveospheres, with or without pre-treatment with TX-100 by ultracentrifugation, compared to unloaded caveospheres. **g**, CryoTEM of SN-caveospheres. Scale bar, 50 nm. **h**, Stability of SN-caveospheres (containing EGFP) after incubation at different pH values at 37°C, followed by neutralizing and purification, compared to untreated SN-caveospheres (Ctrl) using Western blot (1 hr pH treatment). **i**, Loading capacity of sonicated DOX-spheres compared to incubation of DOX and caveospheres without sonication expressed as µg of DOX per 1 mg of caveospheres (n=5 technical replicates over two independent experiments, Student’s t-test). **j**, Fluorescence emission of DOX-spheres after ultracentrifugation assay in the absence or presence of TX-100, compared to unloaded caveospheres (Ctrl). **k**, Negative stained TEM images of unloaded caveospheres and DOX-spheres. Scale bar, 50 nm. **l**, Schematic for generation of SYBR-labeled DNA (SYBR-DNA) loaded caveospheres by sonication (SYBR-DNA-spheres). SYBR-DNA was loaded into caveospheres via sonication followed by DNAse 1 treatment and centrifugal filtration. SYBR-DNA and SYBR-DNA mixed with caveospheres without sonication, treated with DNase I followed by centrifugal filtration were also included. SYBR-DNA without DNase I treatment was used as a control. **m**, Negative stained TEM images of unloaded caveospheres (i) and SYBR-DNA-spheres (ii). Scale bars, 50 nm. **n**, Fluorescence emission of SYBR-DNA-spheres, SYBR-DNA and SYBR-DNA mixed caveospheres (without sonication) after treatment with DNase 1 and purification. **o**, Loading capability (µg DNA in 50 µg caveosphere) of SYBR-DNA loaded caveospheres with and without sonication using fluorescence emission at 522 nm (n=3 technical replicates, Student’s t-test). **p**, Agarose gel electrophoresis of all samples in (n) compared to SYBR-DNA without DNase I treatment. **q**, Fluorescence emission at 525 nm and Western analysis for MBP-Z-domain-CAV1 of fractions collected after sucrose gradient assay of caveospheres + A488 fluorescent antibodybody. Free A488 was used as a negative control. **r**, and **s**, Stability of MBP-Z-domain-CAV1 caveospheres labeled with A488 (A488-sphere) in a biological medium. Ultracentrifugation assay followed by fluorescence imaging of A488-spheres after 0 hrs (no incubation) and 72 hrs incubation in standard tissue culture media (DMEM/10% FBS) (r). Quantification number of A488 antibody (µg) per caveosphere (mg) after 72 hrs at 37°C in DMEM/10% FBS compared to A488-spheres at 0 hrs using the sucrose gradient ultracentrifugation followed by fluorescence emission at 525 nm of fractions at 40% and 60% sucrose levels (s) (t-test, n=3 technical replicate). In all panels: ns: not significant; *: P≤ 0.05; **: P≤ 0.01; ***; P ≤ 0.001****; P≤ 0.0001. The exact P values were provided in supplemental data 10. Error bars represent mean±SD.

In sucrose gradients, mScarlet fluorescence cofractionated with the CAV1 protein as detected by Western blotting within the 40-60% sucrose fractions **(Figure 1c i-ii)**. Cryo-correlative light and transmission electron microscopy (cryoCLEM) of mScarlet loaded GE-caveospheres demonstrated enrichment of the mScarlet fluorescence in areas that correlated with structurally uniform nanovesicles **(Figure 1d)**. Quantitation of the incorporation of mScarlet into caveospheres showed 0.132 ± 0.018 µg mScarlet (mean ± SD) in 1 mg of GE-caveospheres (1.018 ± 0.107 mScarlet molecules per caveosphere) (number of particles per 1mg caveosphere by nanoparticle tracking analysis **Figure S1c**, standard curve **Figure S4a**). Taken together, these results demonstrate that genetically-encoded proteins that are expressed in the periplasm of the bacteria can be encapsulated into the lumen of forming caveospheres, providing a simple system which bypasses the requirement for cargo protein production and purification.

We next tested physical methods of encapsulation which would allow incorporation of diverse cargoes. We evaluated three distinct methods: freeze-thaw cycling, electroporation, and sonication **(Figure S3a)** for encapsulation of EGFP and mScarlet proteins **(Figure 1e & S3c)**. The association between cargo and caveospheres was tested by ultracentrifugation **(Figure 1f & S3d-e)** and negative stain TEM was utilized to evaluate their morphology. All methods demonstrated specific encapsulation of fluorescent proteins while nanovesicle morphology was maintained **(Figure 1f & S3b-e)**. Sonication was the most effective method of encapsulation, exhibiting the highest level of cargo incorporation while retaining the morphology of unsonicated vesicles as judged by cryoelectron microscopy **(Figure 1g)**. The highest loading obtained using this method corresponded to approximately 12 molecules of EGFP per 50nm vesicle (12.2 ± 1.29; **Figure S1c**, standard curve in **Figure S4b**). The use of physical methods, particularly sonication, enables a higher loading capacity compared to genetically encoded approaches, while also broadening the range of cargoes that can be incorporated into caveospheres. In contrast, genetically encoded methods provide a straightforward one-step process for protein loading, making them well-suited for potential scale-up in industrial manufacturing and for the delivery of toxic proteins in cancer therapy, where only small amounts are required to achieve pharmacological efficacy.

Next, we examined the ability of caveospheres to retain their cargo after sonication. EGFP-loaded caveospheres treated at 37°C for 30 min or 1 hour in media with pH from 7.4 to 4.5 retained their contents at neutral pH but released it at lower pH levels **(Figure 1h & S3f)**, revealing an unexpected pH sensitivity.

As a first step in exploring the potential therapeutic applications of caveospheres as an anti-cancer drug delivery vehicle, we loaded caveospheres with Doxorubicin (DOX), a chemotherapeutic agent associated with significant side-effects including cardiotoxicity. ^8–10^ As demonstrated for fluorescent proteins, we were also able to achieve specific encapsulation of DOX while preserving nanovesicle morphology **(Figure 1i-k)**. We next tested the capacity of caveospheres to encapsulate plasmid DNA. Plasmid DNA was fluorescently labeled with SYBR green and loaded into caveospheres via sonication. Any unincorporated plasmid DNA was then removed by DNAse 1 treatment followed by centrifugal filter purification **(Figure 1l)**. Again, negative staining by TEM showed that the morphology of the DNA-loaded caveospheres was maintained **(Figure 1m)**. Encapsulation efficiency was evaluated and quantified using fluorescence. In sonication-loaded spheres, DNA was protected from DNAse1 treatment consistent with encapsulation, whereas DNA incubated in solution with non-sonicated caveospheres was completely cleaved **(Figure 1n-p)**. Based on the fluorescence signal from SYBR Green labelled DNA, we established a linear correlation between SYBR-labeled DNA concentration and fluorescence intensity **(Figure S4d)**. From this analysis, we measured that approximately 2.23 µg of DNA is associated with 50 µg of caveosphere after sonication, corresponding to ∼2.6 plasmid molecules per vesicle **(Figure 1o)**. This level of DNA loading is comparable to that reported for natural extracellular vesicles and outer membrane vesicles (OMVs). ^11–14^

The caveosphere fusion protein contains an IgG binding Z-domain resulting in approximately 150 IgG binding Z-domains per nanovesicle. ^6^ This modularity has the potential to enable targeted delivery to specific cells, tissues and disease states, and allow fluorescent labeling for precise tracking. In order to produce an externally-labelled, trackable particle which retains cargo-loading capacity we specifically bound Alexa-488 conjugated rabbit IgG secondary antibodies to the caveosphere surface via Z-domain. We tested several ratios of fluorescent antibodies to caveospheres and assessed binding using ultracentrifugation. At an antibody-to-caveosphere ratio of 1:50, the fluorescence signal was sufficiently bright to allow reliable detection and tracking of caveospheres **(Figure S4f)**. Under these conditions, we calculated a labeling of 8.87 ± 0.13 IgG molecules per caveosphere (**Figure 1q & S1c**; standard curve shown in **Figure S4e**), corresponding to ∼6% occupancy of the ∼150 theoretical Z-domain epitopes on the caveosphere surface. The use of mouse versus rabbit fluorescent IgG antibodies showed no significant difference in binding levels **(Figure S4g)**. Doubling the antibody ratio increased the surface loading to 13.3 ± 0.73 IgG molecules per caveosphere. Furthermore, co-loading fluorescent IgG and anti-EGFR antibodies (both at 1:50) resulted in 7.2 ± 0.67 molecules of each antibody per caveosphere, with the anti-EGFR level estimated based on fluorescent antibody calibration **(Figure S4g)**. The *in vivo* application of the caveospheres as a potential trackable therapeutic delivery system necessitates stability in complex biological fluids. Fluorescently-labeled caveospheres incubated in tissue culture media with 10% serum at 37°C for 72 hours showed fluorescence quantitatively indistinguishable from the controls **(Figure 1r-s)**, indicating that caveospheres retained their surface antibodies during the incubation period.

### Caveospheres mediate highly specific targeted transfection

We first double-labelled caveospheres with both Alexa-488 antibodies (for tracking) and anti-EGF-R antibodies (for targeting to the human EGF receptor) and validated specific targeting to EGF-R positive A431. Cells in a mixed culture with EGF-negative HepG2 cells **(Figure S5a-b)**. In addition, we loaded caveospheres coated with anti-EGFR with EGFP to test productive delivery into target cells. The cargo was selectively delivered to A431 cells in the mixed culture, with minimal off-target delivery to EGFR-negative cells **(Figure S5c-d).**

Having efficiently incorporated DNA into caveospheres, we next investigated whether targeted caveospheres can deliver DNA to the cytosol and/or nucleus of target cells. EGFR interaction in A431 cells indicates that EGFR-targeted caveospheres were internalized through clathrin-mediated endocytosis. ^15^ Once internalized, caveospheres can fuse with endosomal/lysosomal membranes to release their cargo. ^16–19^ To examine and quantify this process, we employed a DNA construct encoding EGFP, which enabled detection through the appearance of a green fluorescence signal. We loaded caveospheres coated with anti-EGFR with plasmid DNA encoding EGFP and compared EGFP expression to that achieved with a widely used commercial transfection reagent, Lipofectamine, in the same mixed-cell culture system. After incubation in mixed culture specific expression of EGFP occurred exclusively in A431 cells, while HepG2 cells did not show any detectable expression **(Figure 2a-b)**. We term this process caveosphere-mediated targeted transfection (CmTT). In contrast, delivery of the expression vector by Lipofectamine showed no selectivity and EGFP expression was observed in both A431 and HepG2 cells **(Figure 2a-b & S5e)**. Furthermore, the transfection efficiency, expressed as the percentage of cells transfected, was significantly higher with caveospheres (86.9 ± 2.1%) compared to Lipofectamine (47.5 ± 8.1%) in A431 cells **(Figure S5f)**. An additional striking difference between the CmTT method and the Lipofectamine method, was the uniformity of EGFP expression between cells in the CmTT group, in contrast to the Lipofectamine transfected cells where expression was highly variable **(Figure S5g)**. These findings demonstrate two striking advantages of the caveosphere system for transfection; 1) targeted delivery to specific cell types and 2) a greater consistency of expression level over the cell population.

**Figure 2.**
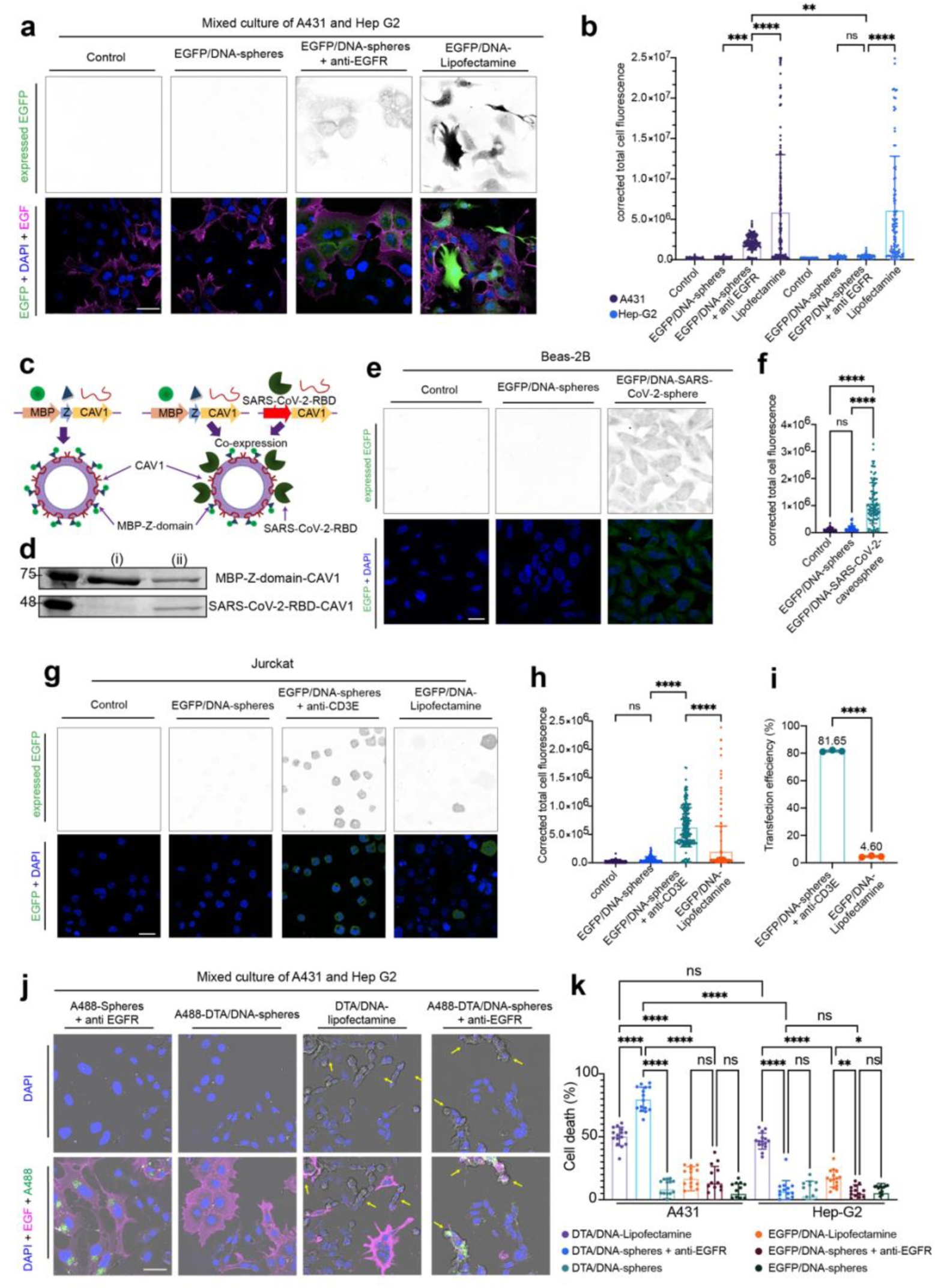
Caveospheres mediate highly specific targeted transfection. **a**, Images of EGFP expression (green) in mixed cultures of A431 and HepG2 cells, stained with Alexa647-conjungated EGF (magenta) and DAPI (blue). Cells were untreated (Control), or transfected with EGFP using either DNA-sonicated spheres expressing EGFP (EGFP/DNA-spheres) with and without anti-EGFR functionalization, or using Lipofectamine (EGFP/DNA-Lipofectamine). EGFP expression also shown as inverted images. Scale bar, 40 µm. **b**, Quantification (corrected total cell fluorescence) of EGFP expression in groups shown in (a) (n=3 independent experiments, one-way ANOVA with Tukey’s multiple comparison test). **c**, Schematic for generation of hybrid SARS-CoV-2-caveospheres co-expressing SARS-CoV-2-RBD-CAV1 and MBP-Z-domain-CAV1. **d**, Coomassie-stained SDS-PAGE gel of purified caveospheres (i) and SARS-CoV-2-caveospheres (ii) with bands corresponding to MBP-Z-domain-CAV1 (72 kDa) and SARS-CoV-2-RBD-CAV1 (44 kDa). **e**, Images of EGFP expression in Beas-2B cells after transfection with EGFP/DNA loaded caveospheres, in comparison to EGFP/DNA loaded SARS-CoV-2-caveospheres and untreated (Control) cells, with DAPI staining (blue). Scale bar, 20 µm. EGFP expression shown as grayscale inverted images. **f**, Quantification (corrected total cell fluorescence) of EGFP expression of groups in (e) (n=3 independent experiments, one-way ANOVA with Tukey’s multiple comparisons test). **g**, Images of EGFP expression (green) in Jurkat cells after transfection with EGFP/DNA loaded caveospheres (EGFP/DNA-sphere +/- anti-CD3E), in comparison to EGFP/DNA-Lipofectamine transfection and untreated (Control) cells, with DAPI staining (blue). Scale bar, 20 µm. EGFP expression also shown as inverted images. **h**, Quantification (corrected total cell fluorescence) of EGFP expression of groups in (g) (n=3 independent experiments, one-way ANOVA with Tukey’s multiple comparisons test). **i**, Comparison of transfection efficiency in Jurkat cells (expressed as percentage of EGFP-positive cells) for EGFP/DNA-sphere + anti-CD3E and EGFP/DNA-Lipofectamine transfection (n=3 independent experiments, student’s T-test). **j**, Images of A431 and Hep G2 after 24 hrs incubation with: caveospheres labeled with A488 and anti-EGFR antibodies (A488-spheres + anti-EGFR), DNA expressing DTA loaded caveospheres labeled with A488 antibodies without or with anti-EGFR functionalization (A488-DTA/DNA-spheres and A488-DTA/DNA-spheres + anti-EGFR, respectively), in comparison to DTA transfection with Lipofectamine (DTA/DNA-Lipofectamine). Cells were stained with Alexa647-conjungated EGF (magenta) and DAPI (blue). Deformed cells are highlighted by yellow arrows. Scale bar, 40 µm. **k**, Cell death (%) of A431 and Hep G2 (in different cultures) after 48 hrs transfection with DTA/DNA-sphere, DTA/DNA-sphere + anti-EGFR or DTA/DNA-Lipofectamine calculated based on cell viability assay. The DNA expressing EGFP was used to detect the effect of protein expression in cell proliferation in three conditions as above (EGFP/DNAsphere, EGFP/DNA-sphere + anti-EGFR and EGFP/DNA-Lipofectamine) (n=3 independent experiments, two-way ANOVA with Tukey’s multiple comparisons test). In all panels: ns: not significant; *: P≤ 0.05; **: P≤ 0.01; ***: P ≤ 0.001; ****: P≤ 0.0001. The exact P values were provided in supplemental data 11 & 12. Error bars represent mean±SD.

mRNA has emerged as a powerful therapeutic modality for vaccination and protein replacement therapy, ^20–22^ motivating our efforts to load mRNA into caveospheres for targeted and efficient intracellular delivery. Functionalized mRNA sonication-loaded caveospheres also gave a transfection efficiency higher than Lipofectamine (% cells transfected), and with less variable expression, similar to the results with encapsulated DNA **(Figure S6 a-c)**.

Next, we explored the potential for decorating caveospheres with genetically encoded binding domains with specificity for cell-surface ligands. We leveraged the high affinity between the receptor-binding domain (RBD) of SARS-CoV-2 virus and the human ACE2 receptor to generate a virus-like particle capable of being selectively endocytosed by bronchial epithelial cells. To this end, we generated a vector encoding which, in addition to the caveosphere fusion protein (MBP-Z-CAV1), also expresses the RBD fused to human Caveolin1 (SARS-CoV-2-RBD-CAV1). This co-expression system generated hybrid spheres containing both the Z-domain and SARS-CoV-2-RBD on the surface **(Figure 2c-d & S6d)**. We expect that the level of RBD displayed on the caveosphere surface correlates with the amount of Caveolin-1 incorporated into the vesicle membrane which is expected to be approximately half to be fused with RBD (at ∼75 RBD per caveosphere).

SARS-CoV2-caveospheres labelled with Alexa-488 antibodies were added to cultures of the human bronchial epithelial cell line BEAS-2B **(Figure S6e-f)**. After loading SARS-Cov2-caveospheres with EGFP plasmid, uptake by BEAS-2B cells resulted in uniform expression of EGFP across the culture **(Figure 2e-f)**. This demonstrates the ability to transduce target cells using diverse receptor-ligand systems and the potential of the caveosphere system for rapid testing of viral protein-host cell interactions.

The antibody-dependent transfection mediated by the CmTT method offers new possibilities for protein expression in cells such as T-cells that have, to date, been refractory to transfection. ^23^ We functionalized caveospheres with anti-CD3E and were able to deliver EGFP into a Jurkat cell line with an efficiency of 81.65 ± 0.6%, significantly higher than Lipofectamine (4.6 ± 0.54%). Conversely, when using non-functionalized spheres, no cells were transfected **(Figure 2g-i)**. This highlights the potential of using caveospheres for delivery to challenging cells, with potential applications in designing and personalizing T-cell therapies. ^24–26^

Finally, we attempted to deliver a DNA cassette with a functional consequence, rather than a benign fluorescent reporter, using caveospheres. We selected DNA encoding diphtheria toxin A (DTA), a highly potent toxin, which must be delivered into the cell cytosol to induce rapid cell death. ^27, 28^ We used CmTT with EGF-R functionalized caveospheres in mixed cultures of A431 and HepG2 cells in comparison to Lipofectamine-mediated delivery. When using Lipofectamine as a delivery method, cell death occurred indiscriminately in the two cell types **(Figure 2k)**. In contrast, the CmTT method resulted in almost complete ablation of A431 cells while leaving HepG2 cells unaffected in the same culture milieu **(Figure 2j and S7)**.

### Tumour targeting and anti-tumour activity of caveospheres in an *in vivo* model

Having demonstrated the stability and targeting capacity of caveospheres in cell culture systems, we investigated their capacity to reach tumor cells *in vivo*. We utilized a well-characterized mouse xenograft model in which an EGFR-positive MDA-MB-468 orthotopic breast cancer cell line is introduced in the left mammary fat pad of a Balb/c nu/nu mouse at 12 weeks of age **(Figure 3a)**.^29^ Caveospheres (200 μg) functionalized with anti-EGFR and Alexa 680 secondary antibody (near-infrared fluorophores used for deeper tissue penetration) were injected intravenously. ^30–32^ After 24 hours, whole animal imaging revealed the specific accumulation of Alexa 680 in the region of the tumor **(Figure 3b)**. Some degree of biodistribution in other organs beyond the tumor was observed due to the intravenous injection. Subsequent imaging of the tumor and specific organs *ex vivo* indicated a significantly higher average radiant efficiency in the tumors from the cohort administered with the anti-EGFR-functionalized caveospheres compared to the cohort given the untargeted spheres **(Figure 3c)**. In contrast, organs such as the liver, lungs, and kidneys exhibited similar levels of caveosphere distribution in both groups. Notably, the heart showed the lowest accumulation. Western blot and immunofluorescence staining against CAV1 in histological sections of tumors showed significantly higher levels of caveosphere-derived CAV1 in the targeted caveospheres cohort than the untargeted cohort **(Figure 3d-f)**. Taken together, these results suggest that antibody labeling of fluorescent caveospheres allows specific targeting to tumors *in vivo*.

**Figure 3.**
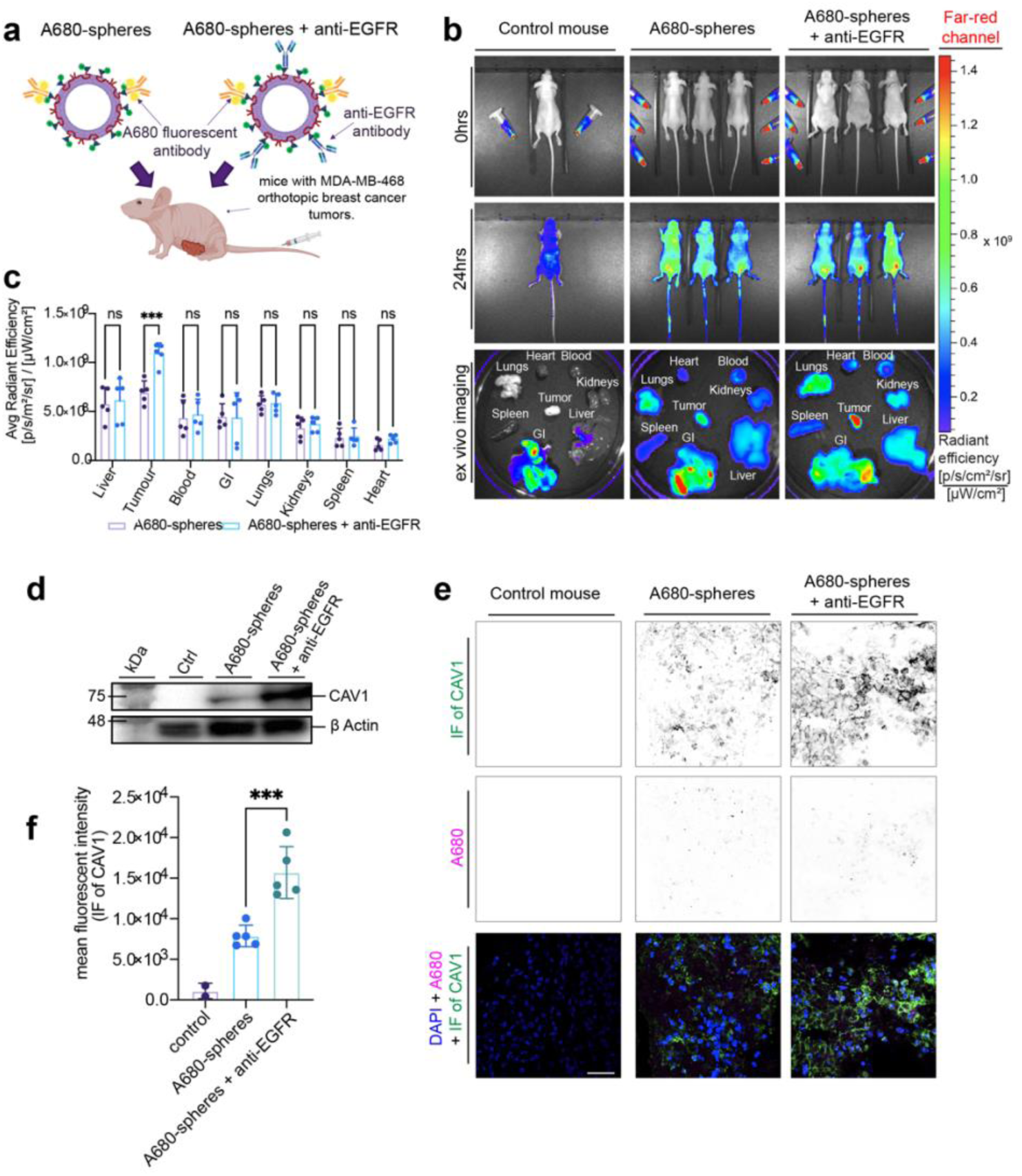
Fluorescent antibody labelling caveosphere for in vivo trafficking and tumour targeting study in mouse xenograft model. **a**, Schematic for mouse xenograft model of orthotopic breast cancer using caveospheres labeled with Alexa-680 secondary antibody (A680-spheres), and functionalized with anti-EGFR antibody for targeting to the MDA-MB-468 derived xenograft. **b**, In vivo imaging of mice injected with control (PBS 1x), A680-spheres or anti-EGFR functionalized A680-spheres (A680-spheres + anti-EGFR). A680 fluorescence shown pre-injection (0 hrs) and 24 hrs post-injection as normalized mean intensity (radiant efficiency) of the near-IR fluorescence is indicated using a blue (low) to red (high) look-up table (n=5 mice per group). Ex vivo imaging of A680 fluorescence of tumor, blood, lung, liver, spleen, kidney, gastrointestinal tract (GI) and heart tissues was also performed (one representative mouse shown). **c**, Quantification of average radiant efficiency in the whole tissue regions of all tissues in both A680-spheres and A680-spheres + anti-EGFR cohorts (after subtraction of values from control mice) (n=5 mice per group, two-way ANOVA with Šidák’s multiple comparison test). **d**, Western analysis of MBP-Z-domain-CAV1 presented in control, A680-spheres, and A680-spheres + anti-EGFR mouse tumor tissue. β-actin is shown as a loading control. **e**, Images of A680 fluorescence (magenta, inverted images in middle panel) from control, A680-spheres, and A680-spheres + anti-EGFR mouse tumor cryosections, stained with anti-CAV1 antibody (green, inverted images in top panel) and DAPI (blue). Scale bar, 40 µm. **f**, Mean fluorescence intensity of CAV1 in tumor cryosections from control (n=2), A680-spheres (n=5) and A680-spheres + anti-EGFR (n=5) mice (Student’s t-test). Dots represent mean intensity of CAV1 imunofluorescence from from 3 randomly selected areas presented in (e). In all panels: ns: not significant; *: P≤ 0.05; **: P≤ 0.01; ***; P ≤ 0.001; ****: P≤ 0.0001.The exact P values were provided in supplemental data 13. Error bars represent mean±SD.

Next, we aimed to test the specificity and efficacy of DOX-loaded caveospheres in the mouse xenograft model. We first tested the DOX-loaded anti-EGFR antibody decorated caveospheres in a cell culture model and showed that they were effective in tumour cell killing as compared to Free-DOX or DOX-loaded caveospheres **(Figure S8a & 4a-b)**. BALB/c nude mice with MDA-MB-468 orthotopic breast cancer were then injected with the anti-EGFR-functionalized DOX-loaded caveospheres, EGFR-functionalized unloaded caveospheres, DOX-loaded caveospheres (without EGFR antibody), Free-DOX, (in the same DOX dose of 4 mg/kg, in 200 µL PBS 1x), and saline **(Figure 4c)**. The general health status of the mice was monitored, and no significant differences were observed between the groups receiving caveospheres with or without anti-EGFR functionalization and the saline-treated control group during the 3-week treatment period (weights given in **Figure S8b**). We observed a significant decrease in the final tumor volume in mice treated with anti-EGFR-functionalized DOX-loaded caveospheres (48.642 ± 11.1 % of the initial volume) **(Figure 4d)**, and more effective tumor growth inhibition than in mice treated with Free-DOX or DOX-loaded caveospheres (without EGFR-functionalization) (Figure 4e-f).

**Figure 4.**
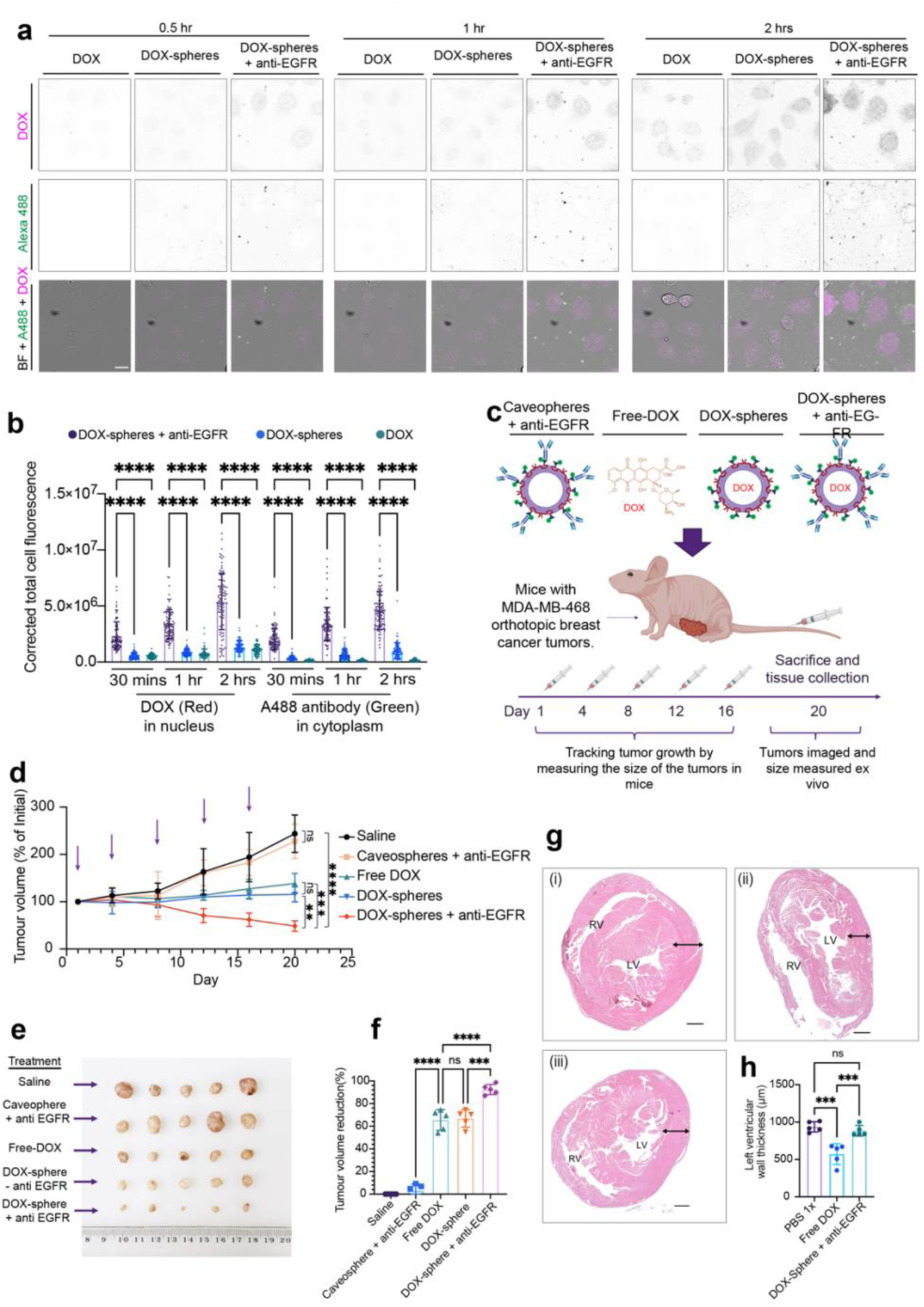
Anti-tumour activity of caveospheres in an in vivo model. **a**, Live-cell confocal imaging of DOX (magenta) and A488 (green) in A431 cells after incubation with free DOX, A488-DOX-spheres or A488-DOX-spheres+anti-EGFR for 0.5, 1 and 2 hrs. BF, Brightfield. Scale bar, 10 µm. **b**, Quantification (corrected total cell fluorescence) of DOX in nucleus and A488 fluorescence in cytoplasm from live-cell confocal imaging in (a) above (n=4 independent experiments, one-way ANOVA with Tukey’s multiple comparisons test). **c**, Schematic of mouse xenograft model of orthotopic breast cancer: Mice were injected with Free DOX, DOX-spheres or DOX-spheres + anti-EGFR on days 1, 4, 8, 12 and 16 of treatment. Saline and anti-EGFR functionalized caveospheres (-DOX) were used as negative controls. **d**, Tumor volumes (expressed a percentage of volume at day 1). Arrows represent days on which mice were dosed. **e**, Excised tumors from mice in (d) after 20 days of treatment. **f**, Tumor inhibition rate (expressed as percentage of reduction in volume compared with saline) in excised tumors after 20 days of treatment. **g**, Representative midventricular cross-section of murine heart from mice in (d) after injection with saline (i), Free DOX (ii) or DOX-spheres + anti-EGFR (iii). Left (LV) and right (RV) ventricles are indicated. Black arrow denotes measurements taken along left myocardium. Tissue was stained with haematoxylin and eosin. Scale bar, 100 μm. **h**, Quantification of left ventricular width from images in (g) (n = 5 mice per group, one-way ANOVA with Tukey’s multiple comparison test). For i-h: n=5 mice per group, one-way ANOVA with Tukey’s multiple comparisons test. In all panels: ns: not significant; *: P≤ 0.05; **: P≤ 0.01; ***: P ≤ 0.001; ****: P≤ 0.0001. The exact P values were provided in supplemental data 14 & 15. Error bars represent mean±SD.

Hearts from experiment above were dissected, embedded in paraffin, stained by hematoxylin/eosin followed by 10 µm histological sections. Quantification of left-ventricular cardiac wall thickness from histological sections as a measure of cardiotoxicity demonstrated a significant decrease in the free-DOX treated mice compared to those treated with anti-EGFR functionalized DOX loaded spheres and saline only controls (Figure 4g-h) demonstrating that targeted delivery encapsulated within caveospheres significantly decreased toxic side effects of the administered DOX.

These data demonstrate that DOX-loaded caveospheres can effectively and rapidly enter both cells and solid tumors to deliver their payload, reducing tumor size and minimizing cardiac side-effects.

### Preparation and characterization of endotoxin-free caveospheres

As a step towards future clinical translation, caveospheres have been engineered to ensure endotoxin levels remain within FDA-approved limits. CAV1 has been expressed in cell-free systems using *Leishmania tarentolae* extract and in Sf21 insect cells to form caveolae with diameters ranging from 50–100 nm. ^33, 34^ However, purification in these systems proved inefficient, resulting in very low vesicle yields, and the capacity for functional domain modification for targeting was limited. Consequently, although mammalian cell–derived systems are endotoxin-free, they are not practical for scalable caveosphere production. ClearColi, a genetically modified E. coli strain lacking endotoxins, has therefore emerged as a promising candidate to address these limitations. ^35^ ClearColi carries seven genetic deletions that prevent the conversion of the precursor lipid IV_A_ into mature lipopolysaccharide eliminating the oligosaccharide chain (including the O-antigen) and two of the six acyl chains that are recognized by the TLR4–MD2 complex **(Figure 5a)**. ^35^ This renders it incapable of triggering an endotoxic response. ^35–37^

**Figure 5:**
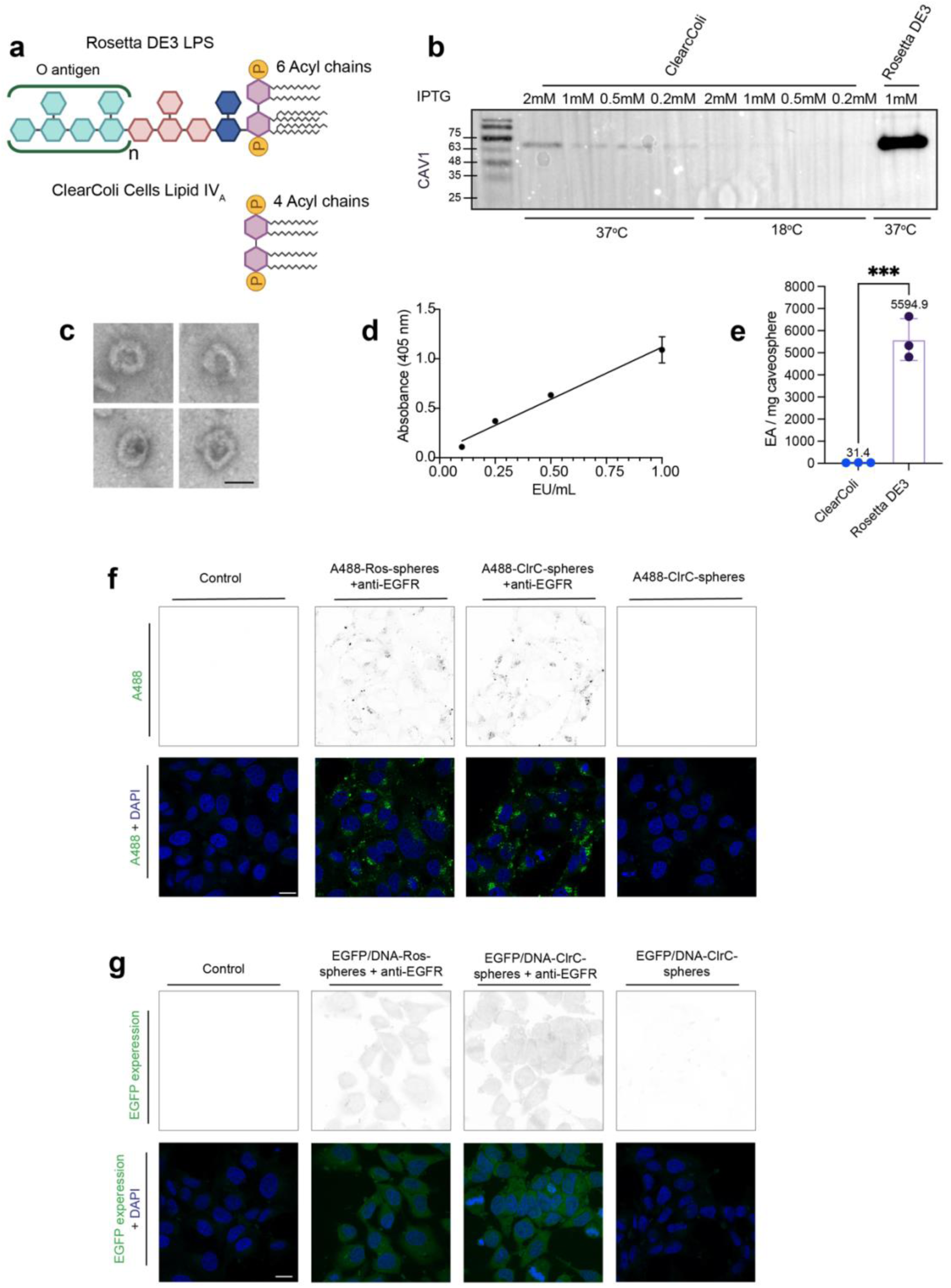
Preparation and characterization of endotoxin-free caveospheres. **a**, Schematic of Rosetta LPS compared to a lipid IVA replacement from ClearColi. **b**, Western blot analysis of MBP-Z-domain-CAV1 (72 kDa) using an anti-CAV1 antibody in 1 mL ClearColi cultures expressing caveospheres after overnight incubation at 18°C or 37°C, induced with 0.2–2 mM IPTG, compared to standard caveosphere expression in Rosetta culture. **c**, Negative-stained TEM image of caveospheres collected from ClearColi culture after purification by column chromatography. Scale bar, 50 nm. **d**, Linear correlation analysis of endotoxin units (EU) measured by Limulus amebocyte lysate (LAL) assay based on absorbance at 405 nm (n = 3 technical replicates). **e**, EU per mg of caveospheres collected from ClearColi and Rosetta, quantified using the LAL assay and the standard curve from (d) (n = 3 technical replicates; Student’s t-test). **f**, Images of A431 cells after 1-hour uptake of caveospheres from ClearColi and Rosetta (ClrC-sphere and Ros-sphere), with or without anti-EGFR targeting antibody. All spheres were labeled with Alexa Fluor 488 (green), and cell nuclei were stained with DAPI (blue). Alexa 488 fluorescence is also shown as inverted grayscale images. Scale bar, 20 µm. **g**, Images of EGFP expression (green) in A431 cells after transfection using ClrC-sphere and Ros-sphere loaded with EGFP plasmid DNA. and cell nuclei were stained with DAPI (blue). EGFP fluorescence is also shown as inverted grayscale images. Scale bar, 20 µm. The exact P values were provided in supplemental data 16. In all panels: Error bars represent mean±SD.

Western blotting against CAV1 confirmed the expression of MBP-Z domain-CAV1 (72 kDa) in ClearColi, and TEM showed that ClearColi-derived caveospheres had a similar morphology and size to those produced in Rosetta DE3 **(Figure 5b-c)**. Although the Limulus Amebocyte Lysate (LAL) assay is the primary FDA-approved method to detect endotoxins, it does not discriminate between endotoxically active hexa-acylated LPS and immunogically inert tetra-acylated lipid IV_A_ in ClearColi. ^35, 38^ Since both share the LAL-activating 4′-monophosphoryl-diglucosamine backbone with acyl chains, ^39, 40^ the LAL assay overestimates endotoxin levels in ClearColi derived products. ^35, 38^ However, caveospheres from ClearColi still showed a drastic reduction in LAL response (∼31 EU/mg vs. >5000 EU/mg from Rosetta), indicating a >99% reduction and allowing for dose optimization to meet FDA safety thresholds **(Figure 5d-e)**. Importantly, these endotoxin-free caveospheres maintained their ability to target cells and deliver plasmid DNA, bringing this system closer to potential use in human therapeutics **(Figure 5f-g)**.

## Conclusion

In this study, we have characterized a unique, modular nanoparticle vesicle as a versatile platform for targeted delivery of encapsulated cargoes. Unlike conventional lipid nanoparticles (LNPs), liposomes or exosomes our system enables straightforward surface modification through simple antibody conjugation via Z-domain or genetically encoded peptide-binding domains, allowing precise targeting to virtually any cell surface antigen. This modularity confers superior targeting and delivery capabilities: only decorated caveospheres—unlike their undecorated counterparts— efficiently deliver diverse cargoes, including DNA, mRNA, proteins, and small-molecule drugs, directly into the cytosol and nucleus of target cells. In comparison with OMVs for delivery applications, caveospheres possess a more versatile surface functionalization module via the genetic fusion of targeting ligands with CAV1 or IgG-binding Z-domain and form vesicles with greater structural homogeneity while most bacterial proteins are excluded during vesicle formation. ^5, 6^ In contrast, OMVs are heterogeneous in size, composition, cargo and their surface proteins can be transferred to host cell membranes, eliciting strong immune stimulation that is advantageous for vaccine development but presents challenges for therapeutic applications. ^41–46^ With lower LPS contamination and more controlled design, caveospheres represent a safer and more versatile alternative to OMVs for therapeutic use. This highlights the caveosphere’s broad utility and potential advantages over existing nanoparticle systems for therapeutic, diagnostic applications, and fundamental biology study.

The power of the CmTT system is illustrated by functional delivery to only the target cells of interest in a mixed culture system (illustrated by anti-EGFR antibodies for tumor cell targeting), and to cells that are difficult to transduce through other means, such as T-cells targeted through surface CD3E. This provides a potential approach to effectively reprogram patient T-cells *ex vivo* in cancer immunotherapy. The ability of CmTT to target specific cell types for efficient delivery of diverse cargoes has great potential for advancing gene therapy applications. The versatility of the system is demonstrated by utilizing the surface Z-domain not only for targeting purposes but also for specific binding to fluorescent antibodies, enabling simple labeling and tracking of nanoparticles *in vitro* and *in vivo*. The caveosphere can be further engineered to mimic virus particles, with the spike protein of SARS-Cov-2 being used as a proof of principle. The establishment of a simple and rapid system for generation of surrogate viruses that effectively enter human target cells offers potential for early-stage screening of antibodies and other molecules to inhibit virus infection. Furthermore, this ability will enable the rapid incorporation of nanobody therapeutics as new targeting sequences are discovered.

Functional delivery of mRNA, DNA, and proteins specifically to target cells with minimal off-target effects is a powerful feature of the caveosphere system. We have demonstrated that this selectivity translates into precision targeting of tumor cells *in vivo*. The high specificity of caveospheres in delivering highly toxic molecules to individual target cells while exerting minimal impact on neighboring cells was demonstrated by a reduction of the adverse effects on the heart compared to the non-packaged therapeutic agent.

The use of ClearColi for caveosphere production does not compromise function and the elimination of LPS is a step towards future use in therapeutics. Although the yield of caveospheres from ClearColi is considerably lower than from Rosetta strains, future studies will address scale-up by optimizing culture conditions and/or the addition of purified individual components to improve vesicle production. The delivery of DNA or mRNA therapeutics in vivo is a promising research direction, however it will require optimization in preclinical models to balance delivery efficiency, and potential side effects, particularly immune activation. ^47–49^ Extended histological analyses of liver and other major organs will also be further studied to avoid unexpected distribution patterns and off-target expression. Beyond applications in diagnosis and therapy, caveospheres loaded with fluorescent proteins and other cargoes can be used to investigate cellular trafficking and endosomal escape of exogenously delivered macromolecules, including screening for genes and/or chemical agents that modulate this process. There is also potential for further manipulation of the caveosphere surface to improve targeting efficiency such as combining surface antibodies with tumor microenvironment–responsive ligands and modifying the physicochemical properties of the vesicles (e.g., surface charge, PEGylation) to extend circulation half-life.

These findings highlight the promise of caveospheres as a simple and rapidly modified system for highly effective transfection in cultured cells, a targeted delivery vehicle for chemotherapeutics *in vivo*, and a platform for imaging and diagnostic applications. The effectively eliminating endotoxin contamination from bacterial sources of the ClearColi caveospheres, supporting their potential for future use in human therapeutics

## Methods

### Cell lines and Culture Conditions

A431 (epidermoid carcinoma, CRL-1555), HepG2 (liver cancer, HB-8065), BEAS-2B (bronchial epithelium, CRL-3588), and Jurkat (T lymphocyte, TIB-152) cells were cultured in their respective media: Dulbecco’s Modified Eagle Medium (DMEM, Cat no. 11965092, ThermoFisher Scientific) for A431 and HepG2 cells, a 50:50 mixture of DMEM and Ham’s F-12 Nutrient Mix (Cat no. 11765054, ThermoFisher Scientific) for BEAS-2B cells, and RPMI 1640 (Cat no. 11875093, ThermoFisher Scientific) for Jurkat cells. All media were supplemented with 10% fetal bovine serum (FBS, Hyclone characterized serum, Quantum Scientific, Lot no. KPJ22093) and L-glutamine (200 mM, Cat no. 25030081, ThermoFisher Scientific). The cells were maintained under standard conditions at 37°C in a 5% CO_2_ incubator. All cell lines were routinely tested for mycoplasma.

### Antibodies

Primary antibodies used in this study were: mouse anti-EGFR antibody (clone LA 22 Merck), mouse Anti-Actin clone C4 (MAB1501) (Merck), Rabbit anti-GFP antibody (A-6455) (Thermo Fisher Scientific), rabbit anti-Cav1 polyclonal antibody (610060) (BD bioscience), Mouse anti-CD3 epsilon antibody (NBP2-53387) (Novus Biologicals).

Secondary antibody used in this study was: rabbit anti-goat IgG secondary antibody, Alexa 488 (Thermo fisher scientific).

### SDS PAGE and Western blot analysis

For SDS-PAGE, caveosphere fractions were denatured with NUPAGE 4x LDS Sample Buffer (Cat no. NP0008, ThermoFisher Scientific) with 10% beta-mercaptoethanol at 95°C for 10 minutes. Proteins were separated on SDS-PAGE gels using Tris-Glycine buffer and were either subjected to Coomassie Blue staining or transferred to an Immobilon-P 0.45 μm PVDF membrane (Cat no. IPVH00010, Merck). The membrane was blocked with 5% skim milk blocking buffer for 1 hour and incubated with a primary antibody overnight. The membrane was then washed with PBST for 10 min three times and incubated with a corresponding secondary antibody (1:5000). The membrane was then washed again with PBST for 10 min three times before visualization of protein bands. Primary antibodies used in this study were rabbit anti-CAV1 (Cat no. 610060, BD Biosciences, 1:5000 dilution), mouse anti-EGFP (Cat no. 11814460001, Roche Diagnostics, 1:5000 dilution), and mouse anti-Actin (Cat no. MAB1501, Merck, 1:5000 dilution) as a loading control. Secondary antibodies for western blotting included Goat anti-Rabbit IgG (H+L) cross-adsorbed secondary antibody, HRP (Cat no. G-21234, Life Technologies, 1:5000 dilution) for rabbit anti-CAV1, and Goat anti-Mouse IgG (H+L) cross-adsorbed secondary antibody, HRP (Cat no. G-21040, Life Technologies, 1:5000 dilution) for anti-EGFP and anti-Actin. Bound IgG was visualized using the Clarity™ Western ECL Substrate (Cat no. 1705061, Bio-Rad). Chemiluminescence was detected using ChemiDoc Imaging System (BIO-RAD). The final images were conducted by merging the chemiluminescence image (showing the WB band) and the colorimetric image (showing the ladder).

Tumor tissue was homogenized using an IKA T10 basic Ultra-Turrax homogenizer. Both tissue and cells were lysed in RIPA buffer containing 50 mM Tris pH 7.5, 150 mM NaCl, 5 mM EDTA pH 8.0, and 1% Triton X-100, supplemented with cOmplete™ mini EDTA-free protease inhibitor cocktail (Cat no. 11836170001, Sigma Aldrich). The lysates were clarified by centrifugation (17,000*g*) at 4°C. Protein content in the tumor samples was quantified using the Pierce BCA Protein Assay Kit (Cat no. 23225, ThermoFisher Scientific) with bovine serum albumin (BSA) as the standard. Forty micrograms of cellular protein were resolved by 10% SDS-PAGE and transferred to an Immobilon-P 0.45 μm PVDF membrane (Merck). Bound IgG was visualized using horseradish peroxidase-conjugated secondary antibodies and Clarity™ Western ECL Substrate (Cat no. 1705061, Bio-Rad).

### Recombinant DNA

Plasmids used in this study were:

– pNMTMA_MBP-Z-Cav1-His6 (Addgene 223701): to produce **standard caveospheres** containing the anti-IgG Z-domain of protein A. Those caveospheres were used for physical loading cargo, cellular uptake, cell transfection and *in vivo* experiments
– pNMTMA_MBP-Cav1-His6 (Addgene 223700): to produce caveospheres minus Z-domain (without the Z-domain on surface)
– pNMTMA_MBP-Cav1-H6_Spike_RBD_Cav1 (Addgene 223702): to produce caveospheres decorated with the spike protein from Sars-Cov2 (SARS-Cov-2-caveospheres)
– pNMTMA_MBP_Z_Cav1_H6_Periplasmic_mScarlet (Addgene 223703): to produce caveospheres with periplasmically encapsulated mScarlet (GE-caveospheres)
– pETDuet-mScarlet (Addgene 223704): for production of mScarlet by *E.coli*
– pEGFP-C1 (Clontech): Mammalian expression vector encoding EGFP-C1
– DTA in pCDEST2 (Addgene 223705): Mammalian expression vector encoding Diptheria toxin A
– pOPINE-GFP (Addgene 223706): for production of EGFP by *E.coli*
– pETDuet-mNeonGreen (Addgene 230970): for production of mNeonGreen by *E.coli*
– pETDuet-nls-mNeonGreen (Addgene 230971): for production of mNeonGreen incorporating an N-terminal nuclear localization signal by *E.coli*
– pETDuet-nls-7R-mNeonGreen: (Addgene 230972) for production of mNeonGreen incorporating an N-terminal nucleolus localization signal (NLS + 7 arginine) by *E.coli*

### Caveosphere preparation

Caveospheres were produced as previously described. ^6^ Briefly, *E. coli* (Rosetta DE3) carrying caveosphere-expressing plasmids were initially cultured in LB and transferred to Terrific Broth at 37°C. Recombinant CAV1 fusion protein induction and caveosphere formation were achieved by adding 1 mM IPTG (Astral Scientific) to the bacterial medium. Following overnight culture at 37°C, cells were lysed using a cell disruptor (Constant Systems), and cellular debris was removed via centrifugation (15,000*g*, 30 min). Caveospheres were purified using an amylose resin (NEB) column affinity chromatography based on surface protein MBP.

### Recombinant protein production

mScarlet, EGFP, mNeongreen, NLS-mNeongreen and NLS-7R-mNeongreen were produced by expression in *E. coli* (Rosetta DE3) overnight followed by TALON® Superflow™ histidine-tagged column affinity chromatography. The proteins were concentrated using Amicon Ultra Centrifugal Filters-0.5 10 kDa MWCO (Merck Millipore) and frozen at a stock concentration of 8 mg/mL.

### Caveosphere quantification

The quantification of produced caveospheres was based on their protein component concentration in solution, determined using the Pierce™ Bicinchoninic Acid Assay Kit (BCA protein assay) (Thermofisher). To analyze the nanoparticles (NPs), 1 mg/mL of protein caveosphere underwent nanoparticle tracking analysis (NTA) utilizing NanoSight NS300. This analysis determined both the number of particles in 1mg/ml protein caveosphere solution and the hydrodynamic diameter distribution of the particles.

### Imaging of fluorescence within microcentrifuge tubes

Microcentrifuge tubes were imaged using a ChemiDoc MP Imaging System (BIO-RAD; red channel 605/50 filter, green channel 530/28 filter).

### Testing of encapsulation by ultracentrifugation and copper ions

Caveospheres with genetically-encoded mScarlet (GE-caveospheres) were incubated in 1% Triton X-100, 37°C for 1 hour to disrupt the particle membrane. The standard GE-caveospheres and the disrupted GE-caveospheres were centrifuged at 120,000*g* for 2 hours in an TLA-55 Fixed-Angle Rotor of Optima MAX-XP Ultracentrifuge (Beckman coulter) followed by imaging of fluorescence using a ChemiDoc MP Imaging System as above. To determine the effective concentration for fluorescence quenching by copper ions, different concentrations of CuCl_2_ (5-0.156 mM) were incubated with free mScarlet (0.1 mg/mL) for 1 hour at room temperature. Then, 1mg/mL of GE-caveospheres, GE-caveosphere treated with Triton X-100 or free mScarlet (0.2 mg/mL) were then mixed with 0.02 mM CuCl_2_ and incubated for 1 hour at room temperature.

### Sucrose gradient analysis of caveospheres

For sucrose gradient analysis, the caveosphere mixture was layered onto a 20%–80% discontinuous sucrose gradient and subjected to velocity gradient centrifugation (120,000*g*, 6 hours, SW 41 Ti Swinging-Bucket Rotor, Beckman Coulter). Twenty fractions were collected from the bottom of the tube, numbered, and transferred to a 96-well plate for western blotting using rabbit anti-CAV1 antibody (BD Bioscience). To determine the presence of GE-caveospheres, fluorescent measurements of all fractions were conducted using a microplate reader as above.

### Quantification of fluorescent proteins, antibodies, DNA and doxorubicin using a microplate reader

To quantify fluorescent protein concentration, the linear correlation between fluorescent protein concentration and fluorescent signal was measured using a microplate reader. For mScarlet, the concentration range was 0.575 mg/mL - 1.4x10^-4^ mg/mL (λex 569 nm, λem 594 nm). For EGFP, the range was 1.16 mg/mL - 4.5x10^-4^ mg/mL (λex 488 nm, λem 507 nm). Sample concentrations were calculated using the linear regression equation derived from the standard curves provided in the supplementary information.

For quantification of antibody labeling of caveospheres, the same plate reader methodology was used to establish the linear correlation between concentration of fluorescent antibodies and fluorescent signal (λex – 490 nm and λem – 525 nm). The microgram quantity of antibody binding per mg of caveospheres was calculated using the same methodology based on the fluorescence (λex – 490 nm and λem – 525 nm) and the protein concentration of caveospheres determined by a BCA protein assay as above at the fractions 9-16 (40-60% sucrose density). Number of antibody molecules per single caveosphere was calculated according to the following equation:

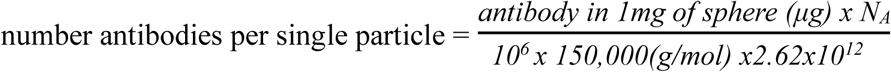

DNA concentration was measured by establishing the linear correlation between SYBR™ Safe labeled DNA concentration and fluorescence (λex 498 nm, λem 522 nm). Doxorubicin concentration was determined in the same way using fluorescence absorbance at 475 nm.

### Electron microscopy of caveospheres

Negative staining was performed by applying 4% uranyl acetate to 12 µL of caveospheres on formvar-coated grids, followed by grid drying and examination under a JEOL 1011 electron microscope at 80kV accelerating voltage. For cryo-electron microscopy, GE-caveospheres were added to R2/2 Quantifoil grids and plunge-frozen using a Leica Electron Microscopy Grid Plunger (Leica Microsystems). Grids were imaged on a CryoARM300 (JEOL) fitted with an in-column Omega filter and K3 camera (Gatan) under low dose conditions. Imaging of EM studies were performed in a blinded fashion.

### Physical encapsulation methods of fluorescent protein into caveospheres

Purified fluorescent proteins were mixed with 1 mg/mL of caveospheres and treated with the following physical methods: freeze-thaw (FT) method (liquid nitrogen -196°C for 5 minutes, followed by thawing in an ice bath for 90 minutes over 4 cycles), electroporation (EL) method (2.5 kV, 75 μF, using the Gene Pulser II electroporation system, BIO-RAD) or sonication (SN) method (20% power with 3 seconds on/6 seconds off for 10 cycles using a Qsonica Sonicator Q700, Fisher Scientific). The incubated caveospheres (IN) in each case was an aliquot of the same sample incubated on ice alongside those undergoing encapsulation. Free mScarlet/EGFP proteins after the treatment process were removed using Amicon Ultra Centrifugal Filters-0.5 100 kDa MWCO (Merck Millipore). The fluorescent signal was then quantitively measured using a microplate reader as caveosphere concentration in the solution was measured by a BCA assay as above.

### Acid treatment of EGFP loaded caveospheres

Sonicated EGFP-caveospheres were incubated in citric buffer at decreasing pH levels (6.0, 5.5, 5.0, 4.5) and in PBS 1x (pH 7.4) for 30 min and 1 hour at 37°C. After incubation, the samples were neutralized with PBS 1x, centrifuged to remove precipitates, and filtered to eliminate fragments. The remaining EGFP-caveospheres were quantified and compared to untreated samples using a BSA assay for caveosphere quantification (as above) and fluorescence measurement with a microplate reader (as above). Western blot analysis was performed against CAV1 using specific antibodies as above.

### Fluorescent antibody-labeling of caveospheres

10 μg standard caveospheres were mixed with 0.2 μg of rabbit anti goat-Alexa 488 in 100 μL of PBS 1x and incubated for 1 hour at 37°C. 800 μL of PBS 1x was added to each tube, followed by ultracentrifugation (2 hours, 120,000*g*) and imaged in microcentrifuge tubes as above. Samples were fractionated by sucrose gradients, and fluorescence was quantified using a microplate reader as above. CAV1 content was analyzed using western blotting as above.

To assess the labeling stability on caveosphere, A488-antibody-labeled caveospheres were incubated in DMEM with 10% FBS, 1 % L-glutamine for 72 hours at 37°C. After 72 hours the stability of A488-caveospheres was compared with freshly prepared A488-caveospheres using ultracentrifugation and sucrose gradients as described above. The amount of Alexa 488 fluorescent antibody on the caveosphere was calculated based on their fluorescent signal (λex – 490 nm and λem – 525 nm) determined by a microplate reader and formula described as above.

### Loading of DNA, mRNA and DOX into caveosphere using sonication

To label plasmid DNA, SYBR™ Safe DNA Gel Stain (Thermo Fisher Scientific) was added to the plasmid DNA at a dilution of 1:10,000. Excess SYBR was removed with an Amicon Ultra Centrifugal Filter-0.5 100 kDa MWCO (Merck Millipore). The concentration of SYBR-DNA was measured using a microplate reader as described above.

For loading of caveospheres, SYBR-labeled plasmid DNA (1000 ng/μL) was mixed with caveospheres (1 mg/ml) and sonicated termed SYBR-DNA-spheres. To remove unloaded DNA, DNAse 1 was added (179 units at 37 °C for 1 hour), followed by filtration. Mixtures without sonication and DNA without caveospheres were used as controls. Samples in microcentrifuge tubes were imaged using a ChemiDoc system as above and analyzed via agarose gel electrophoresis. The extent of DNA encapsulation was quantified by treating SYBR-DNA-spheres with Triton X-100 (1% for 1 hour at 37°C) and was measured using a microplate reader as described earlier. mRNA expressing EGFP (BASE mRNA facility, University of Queensland; 300 ng/μL) was mixed with caveospheres (1 mg/ml) sonicated and purified using sucrose gradient described above.

For doxorubin (DOX) loading, caveospheres (1 mg/mL) were mixed with DOX (0.1-0.5 mg/mL) and sonicated. DOX-caveospheres were treated with Triton X-100 (1% for 1 hour at 37°C) to release DOX, and the loading efficiency was determined using a plate reader as above.

### Functionalization of caveospheres for targeting

Fluorescently-labeled caveospheres or cargoes (EGFP, DNA, mRNA, DOX) loaded caveospheres, were functionalized with or without mouse anti-EGFR clone LA 22 at a mass ratio of 50:1. Unbound antibodies were removed using sucrose gradients following the method described earlier.

### Cellular uptake of caveospheres

For cellular uptake of caveospheres, cells (A431, HepG2, BEAS-2B) were plated on glass coverslips and cultured overnight until reaching a 60-80% confluence. Anti-EGFR-funtionalized caveospheres (containing A488 antibody or EGFP) prepared at a final concentration of 50 μg/mL were incubated with A431, HepG2 and mixed culture of two cell lines on cover slips for 1 hour at 37°C. SARS-CoV-2-caveospheres (labeled with A488) were incubated with BEAS-2B on over slips for 3 hours at 37°C.

### Cell transfection

For cell transfection experiments, caveospheres were loaded with either plasmid DNA encoding EGFP, mRNA encoding EGFP or plasmid DNA encoding diphtheria toxin A (DTA). 150 µg/mL anti-EGFR functionalized DNA/mRNA-caveosphere or DNA-SARS-CoV-2-caveosphere (containing 687 ng DNA or 299 ng of mRNA) was incubated with the cells (A431, HepG2, mixed culture or Beas-2B) on glass coverslips or 6 well plates for 1 hour and washed with standard media. Lipofectamine 2000 transfection of EGFP and DTA was conducted as per the manufacturers protocol. The cells were incubated at standard conditions (at 37°C with 5% CO_2_) for 24 hours.

For Jurkat cell transfections, 10^6^ cells in 500 mL were transfected in a 24 well plate using anti-CD3e functionalized DNA-caveospheres and Lipofectamine as above. An additional spin down protocol using centrifugation (3,000*g*, 5 minutes) was applied for adhesion of cells on coverslips.

After 24 hours, western blotting was performed to evaluate EGFP expression levels in cell lysates collected from each well plate. Expression levels and transfection efficiency were also assessed via confocal microscopy of fixed cell on coverslip described below.

### Confocal imaging of fixed cells and Image analysis

For confocal imaging of fixed cells, cells on coverslip were fixed with 4% paraformaldehyde and stained with DAPI. EGF-conjugated Alexa 647 dye at 2 μg/mL was applied to mixed culture. The coverslips then were imaged using a Zeiss LSM880 Fast Airyscan Confocal Microscope. ImageJ software (NIH) was used for flat-field correction of images using the BasiC plugin ^50^. The green fluorescence intensity of 25-35 randomly imaged cells was determined by calculating corrected total cell fluorescence (CTCF) via the equation; CTCF = IntDen − (Area of selected cells X background mean grey value) using ImageJ, where IntDen is “Integrated Density”. ^51, 52^ From the confocal imaging of fixed cells, total of 90 cells were counted and calculated percentage of EGFP-positive cells for transfection efficiency (%) in A431 cells and Jurkat cell.

### Encapsulated Protein Delivery

50 μg/mL caveospheres loaded with mNG, NLS-mNG, and NLS-7R-mNG plus anti-EGFR antibody were added to A431 cells in an 8-well chamber slide (Nunc™ Lab-Tek™ II CC2™) and incubated at 37°C. Where relevant, chloroquine (100 μM) and bafilomycin A1 (1μM) were added for an hour of incubation. Live cell imaging was conducted on the Zeiss LSM880 Fast Airyscan confocal microscope over 3 hours, capturing 3 random fixed positions in each well at 1, 3-hour time points. ImageJ was used to measure the intensity of green fluorescence in the cell cytoplasm, nucleus and quantified using CTCF quantification in four independent experiments as described above.

### DOX-loaded caveosphere experiments

The DOX, DOX-loaded caveospheres, and DOX-loaded caveospheres plus anti-EGFR antibody at a 2 μg/mL DOX dose were added to A431 cells in an 8-well chamber slide (Nunc™ Lab-Tek™ II CC2™). Live cell imaging was conducted on the Zeiss LSM880 Fast Airyscan confocal microscope over 2 hours, capturing 4 random fixed positions in each well at 0.5-, 1-, and 2-hour time points. ImageJ was used to measure the intensity of green fluorescence in the cell cytoplasm and red fluorescence in nuclei from 15-30 cells at 0.5, 1, and 2 hours, quantified using CTCF quantification in four independent experiments as described above.

### Cell viability test

To assess the viability of cells after DTA transfection by caveospheres/Lipofectamine or DOX treatment, the cells at 48 hours incubation time in 96 well plate were treated with PrestoBlue cell viability reagents for 3 hours followed by fluorescence reading using a plate reader (λex – 560 nm and λem – 590 nm). The cell death percentage was calculated based on the fluorescence (subtract background) of treated cell and untreated cells (100% survival baseline). Cell viability after treatment with chloroquine and bafilomycin A1 was performed as above but after 24 hours.

### *In vivo* mouse experiments

Eight-week-old female Balb/c nude Nu/Nu mice were used for *in vivo* targeting experiments. All animal experiments were approved by the University of Queensland’s Animal Ethics Committee: Anatomical Biosciences AEC (ABS) committee (Project Number: 2019/AE000105) and the Laboratory Biomedicine AEC - LBM committee (Project Number: 2022/AE000135) and conformed to the guidelines of the Australian Code of Practice for the Care and Use of Animals for Scientific Purposes (AEC approval number: AIBN/CAI/530/15). For all animal models, 8-week-old Balb/c nude mice were acquired from the Animal Resource Centre and were allowed access to food and water *ad libitum* throughout the course of the experiment. Tumors were established by injecting 3 × 10^6^ MDA-MB-468 cells into the left mammary fat pad of anesthetized mice. After 12 days of tumor growth, mice with palpable ∼5 mm diameter tumors were selected.

### Tumor targeting

For targeting experiments, caveospheres (200 μg) labeled with Alexa 680 secondary antibody with or without anti-EGFR antibody (mass ratio 50:1) were prepared and injected intravenously into the mice’s tail veins of the mice. Imaging was performed 24 hours post-injection using an IVIS® Lumina™ X5 imaging system. After imaging, mice were humanely sacrificed, and their organs were imaged ex vivo by an IVIS® Lumina™ X5 imaging system.

### Immunofluorescence and western blot of tumor

Tumor samples from above were collected and snap frozen in liquid Nitrogen for Western analysis (as described above), or embedded in O.C.T. Compound (Tissue-Tek®) and frozen in isopentane-cooled liquid nitrogen for histology. For histology analysis, 20 µm cryosections were fixed in 4% PFA, followed by blocking in 2% BSA/PBS 1x. Sections were incubated in primary primary rabbit anti-CAV1 antibody (overnight) and secondary antibodies (1 hour at room temperature) diluted in blocking solution. Images were captured using a Zeiss LSM880 Fast Airyscan Confocal Microscope. For quantification, tissue sections were imaged at three random locations, and mean grey values in the green channel were collected using ImageJ software.

### Tumor inhibition

DOX-loaded caveospheres + anti-EGFR were prepared (DOX dose of 4 mg/kg) and administered systemically to mice along with other treatment groups (free DOX, DOX-caveospheres and anti-EGFR functionalized DOX-caveospheres). Each treatment group consisted of 5 mice with 5 injections over 20 days. Tumor growth was monitored using an electronic digital caliper and calculated using the standard volume formula. ^53, 54^ Then mice were humanely sacrificed on day 20 and the tumors and hearts were collected. ^55–57^ Tumor inhibition rates were calculated on day 20 compared to the control group using this formula:

(1 – (mean volume of treated tumors)/(mean volume of control tumors)) × 100%.

Hearts were dissected, fixed in 4% PFA in PBS 1x at 4 °C for 24 hours. Tissue was embedded in paraffin and routine hematoxylin and eosin staining was performed on 10 µm sections by the Queensland Brain Institute histology service at The University of Queensland. Slides were scanned using a Metafer VSlide Scanner by MetaSystems using Zeiss Axio Imager Z2. Images were processed using the QBI Batch SlideCropper and ImageJ. Left ventricular width was calculated using ImageJ.

### Limulus amebocyte lysates (LAL) assay

The endotoxin content per caveosphere was measured by LAL assay using the Pierce™ Chromogenic Endotoxin Quant Kit (Thermo Fisher Scientific), following the manufacturer’s instructions. A linear standard curve of endotoxin units (EU) was generated to quantify the endotoxin concentration.

Caveospheres produced from E. coli Rosetta and ClearColi strains were first processed through the Pierce™ High-Capacity Endotoxin Removal Spin Column (Thermo Fisher Scientific). After purification, the caveospheres were diluted to final concentrations of 0.15 µg/mL (Rosetta) and 8 µg/mL (ClearColi) prior to performing the LAL assay.

### Software

Figures were prepared using ImageJ and Adobe Illustrator. Schematic images were created with https://BioRender.com and Microsoft office. Statistics were performed using GraphPad Prism9.

### Statistics and replication

Details of all statistical tests, replication and experimental groups are given in the figure legends. In all panels ns: P > 0.05, *: P≤ 0.05, **: P≤ 0.01, ***: P ≤ 0.001****: P≤ 0.0001. The exact P values and statistics were provided in supplemental data 10-16.

### Reagents

Paraformaldehyde (Cat no. P6148-500G, Sigma-Aldrich), Triton X-100 (Cat no. T9284, Sigma-Aldrich), PBS Tablets pH7.2 1000mL/tab.100 tablets (Cat no. 09-9499-10, Astral Scientific), Isopropyl β-D-1-thiogalactopyranoside (IPTG) (Cat no. BIO-37036, Bioline), 4’, 6-diamidino-2-phenylindole (DAPI) (Cat no. D9542-5MG, Sigma-Aldrich), Amylose Resin (Cat no. E8021L, Biolabs), TALON® Superflow™ (Cat no. GEHE28-9575-02, Bio-Strategy), Terrific Broth (Cat no. 22711022, Thermo Fisher Scientific), Ampicillin Sodium (Cat no. 016-23301, Novachem), EGF conjugated Alexa 647 dye (Cat no. E35351, Thermo Fisher Scientific), Doxorubicin hydrochloride injection solution (Cat no. 23214-92-8, Pfizer), SYBR™ Safe DNA Gel Stain (Cat no. Thermo Fisher Scientific), DNAse 1 (Cat no. 18047019, Thermo Fisher Scientific), Lipofectamine 2000 reagent (Cat no. Invitrogen), ECL detection reagent (Life Technologies), BCA protein assay kit (Cat no. 11668019, Thermo Fisher Scientific), Sodium chloride (Cat no. S9888, Sigma-Aldrich), Copper Chloride (Cat no. 222011, Sigma-Aldrich), Sucrose (Cat no. SA030-5KG, Chem-supply), isopentane (Cat no. 270342-1L, Sigma-Aldrich), O.C.T. Compound (Cat no. 4583, Tissue-Tek®). The lysosomotropic agents: Amantadine (A1260), Azithromycin (75199), Chloroquine (C6628), Dimebon (D6196), Tamoxifen (T5648), Amitriptyline (A8404), and Siramesine (SML0976) were from Sigma-Aldrich, while UNC10217938A was from Focus Biosciences (Cat no. HY-136151).

### Inclusion & ethics statement

This research included local researchers throughout the research process – study design, study implementation, data ownership, intellectual property and authorship of publications. This research is locally relevant – this has been determined in collaboration with local partners. Roles and responsibilities were agreed amongst collaborators ahead of the research and capacity-building plans were discussed. This research has not been severely restricted or prohibited in the setting of the researchers. This study has been approved by a local ethics review committee (animal ethics noted above). The research does not result in stigmatization, incrimination, discrimination or otherwise personal risk to participants. Risk management plans were undertaken in according to University of Queensland guidelines. Benefit sharing measures have been discussed in case biological materials have been transferred out of the country. Local and regional research relevant to this study has been taken into account in citations.

## ASSOCIATED CONTENT

## Supporting Information

Figure S1-S9 provide supporting data for Figure 1-5 in the main text.

Figure S10-S16 provide exact P values and statistics for Figure 1-5 in the main text.

## Data and materials availability

The data that support the findings of this study are available from the corresponding author upon reasonable request.

## Author Contributions

Conceptualization, R.G.P., T.D.L., A.P.R.J., T.E.H.; methodology, T.D.L., R.G.P., T.E.H., N.M., J.R., N.A., A.P.R.J., N.F; formal analysis, T.D.L., R.G.P., T.E.H., N.M., J.R., H.S., N.A., A.P.R.J., N.F; data curation, T.D.L., Y.-W.L.; funding acquisition, R.G.P., A.P.R.J.; supervision, R.G.P., T.E.H., A.P.R.J, K.T.; writing – original draft, R.G.P., T.E.H., T.D.L., H.P.L, Y.W., K.A.M., Y. - W.L.; writing – review & editing, R.G.P., T.E.H., T.D.L., H.P.L, Y.W., K.A.M., Y.-W.L., A.P.R.J., N.F., K.T.; project administration, R.G.P., T.E.H., A.P.R.J.

## Funding Sources

The National Health and Medical Research Council of Australia grants APP1140064 and APP1150083 to R.G.P.

The Australian Research Council (Centre of Excellence in Convergent Bio-Nano Science and Technology CE140100036 to R.G.P and A.P.R.J).

NHMRC fellowship APP1156489 and is now an Australian Research Council (ARC) Laureate Fellow to R.G.P.

Human Frontiers Science Program Grant (RGP011/2023) to N.A.

## Declaration of interests

The authors declare no competing interests

## ACKNOWLEDGMENT

This work was supported by the National Health and Medical Research Council of Australia grants APP1140064 and APP1150083 to R.G.P. and by the Australian Research Council (Centre of Excellence in Convergent Bio-Nano Science and Technology CE140100036 to R.G.P and A.P.R.J). RGP was supported by an NHMRC fellowship APP1156489 and is now an Australian Research Council (ARC) Laureate Fellow. N.A is supported by a Human Frontiers Science Program Grant (RGP011/2023). The authors acknowledge the use of the Microscopy Australia Research Facility at the Centre for Microscopy and Microanalysis at The University of Queensland.

## ABBREVIATIONS

TEM: Transmission electron microscopy
DOX: Doxorubicin
CmTT: caveosphere-mediated targeted transfection
RBD: receptor-binding domain
DTA: diphtheria toxin A
LV and RV: Left and right ventricles
LPS: Lipopolysaccharide
LAL: Limulus Amebocyte Lysate.

## Notes

### Competing Interest Statement

The authors have declared no competing interest.

### Summary of Updates

This version of the manuscript was added with the preparation and characterization of endotoxin-free caveospheres in Figure 5. These endotoxin-free caveospheres maintained their ability to target cells and deliver plasmid DNA while reducing the endotoxin to meet FDA safety thresholds, bringing this system closer to potential use in human therapeutics. The section on endosomal escape was removed to strengthen the manuscript direction toward the precise delivery of caveosphere in vitro and in vivo.

