## Supplemental data for "A modular encapsulation system for precision delivery of proteins, nucleic acids and small molecules"

---

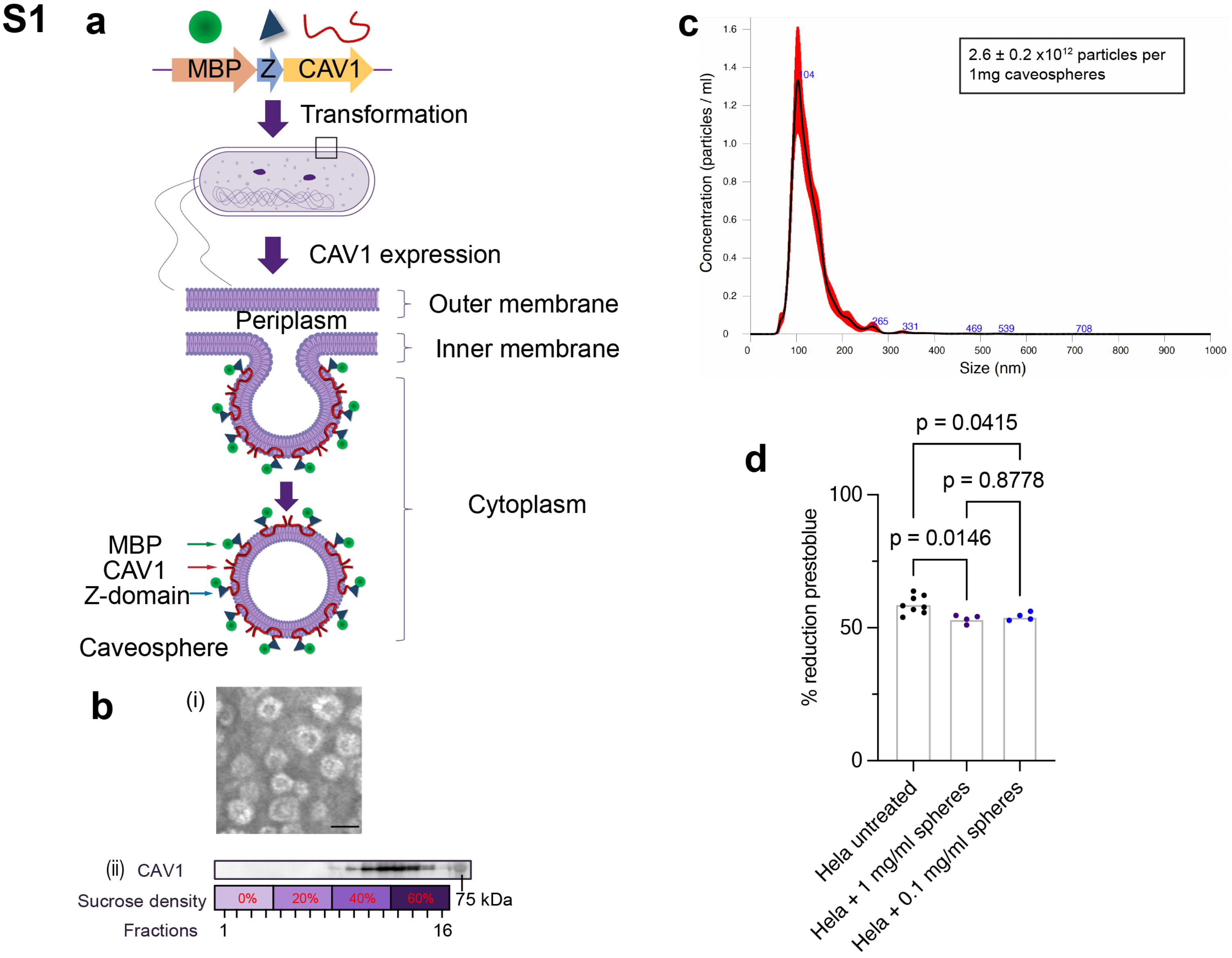

**Supplementary Figure 1: a,** Caveospheres were formed in E. coli by expression of a fusion protein consisting of Canis lupus familiaris Caveolin1 (CAV1), Maltose-binding protein (MBP) and the Z-domain of protein A (to allow IgG conjugation). Those caveospheres were set as standared caveospheres and used for physical loading cargo, cellular uptake, cell transfection and in vivo experiments. **b,** (i) Transmission electron microscopy (TEM) image of caveospheres. Scale bar, 50 nm. (ii) Western analysis for MBP-Z-domain-CAV1 (72kDa) using anti-CAV1 antibody in fractions collected after a discontinuous sucrose gradient assay of caveospheres (0%, 20%, 40%, and 60%, 4 fractions per concentration). Molecular weight marker (75kDa) as indicated. **c,** Nanoparticle tracking analysis report of caveospheres: Finite track length adjustment (FTLA) analysis shows an average caveosphere diameter of 107 ± 4 nm (n = 5 technical replicates, mean - black line, red area - ± SD). Concentration measurements provide an average of particles per 1 mg caveospheres (2 independent experiments, 5 technical replicates per experiment, ± SD). **d,** Prestoblue Cell Viability assays for cytotoxicity testing of Hela cell treated with caveosphere (0.1 and 1 mg/mL) for 24hrs (4 technical replicates per experiment, ± SD).

S2

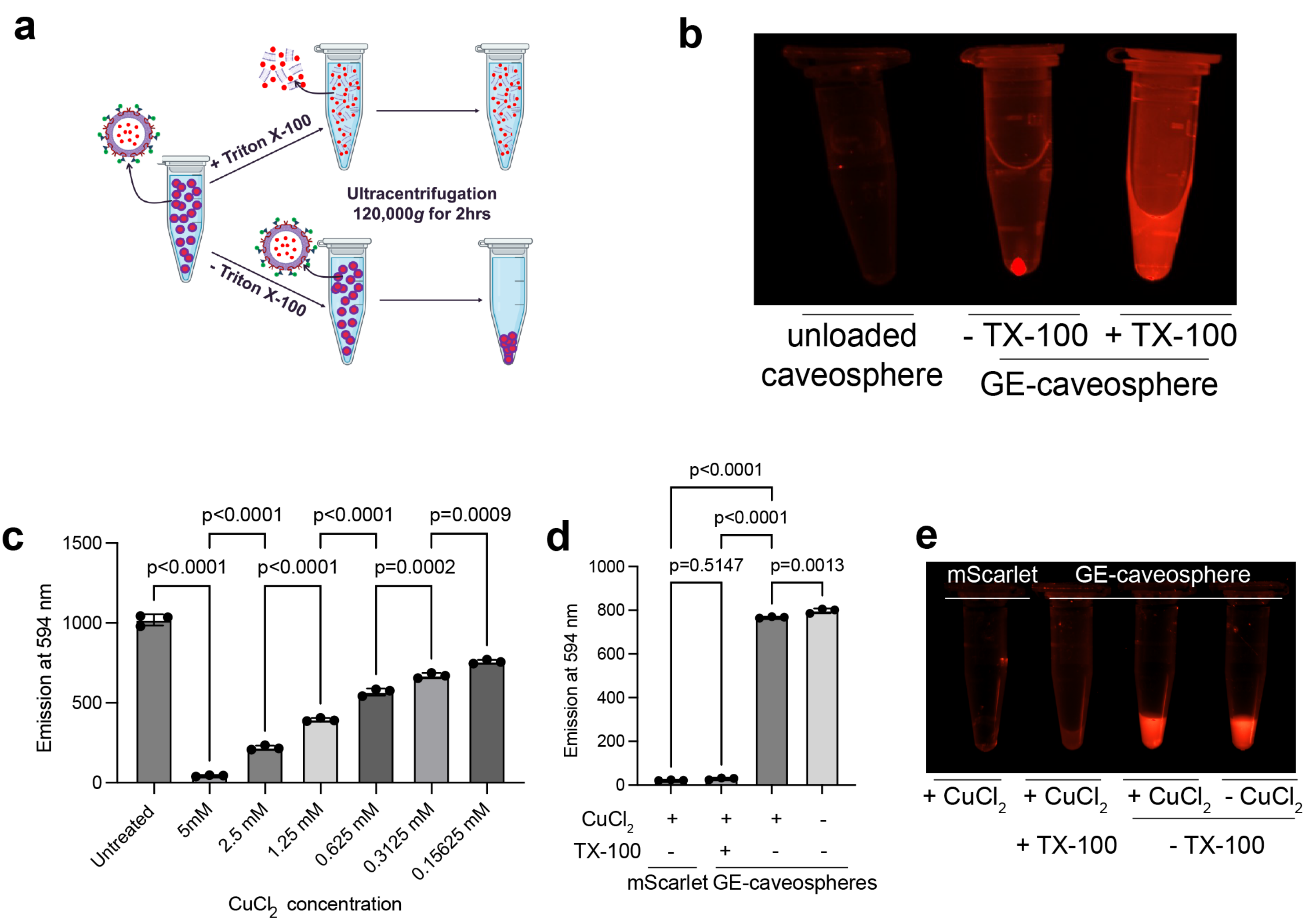

**Supplementary Figure 2: a,** and **b,** mScarlet loaded GE-Caveospheres are pelleted by ultracentrifugation at 120,000g. Pre-treatment with TritonX-100 (TX-100) detergent to disrupt the caveosphere membrane prevents pelleting. Fluorescence emission of mScarlet encapsulated in GE-caveospheres was then captured using a fluorescent imaging system in (b). **c,** Quantification of mScarlet fluorescence intensity at 594 nm at different $CuCl_2$ concentrations. Concentration of mScarlet protein was 0.1 mg/mL (n=3 technical replicates, one-way ANOVA, Tukey's multiple comparisons test). **d,** Fluorescence intensity at 594 nm of free mScarlet and mScarlet-GE-caveospheres treated with $CuCl_2$ and TX-100 (n=3 technical replicates, one-way ANOVA, Tukey's multiple comparisons test). **e,** Fluorescence emission of free mScarlet and mScarlet-GE-caveospheres treated with $CuCl_2$ and TX-100. In all panels: Error bars represent mean±SD.

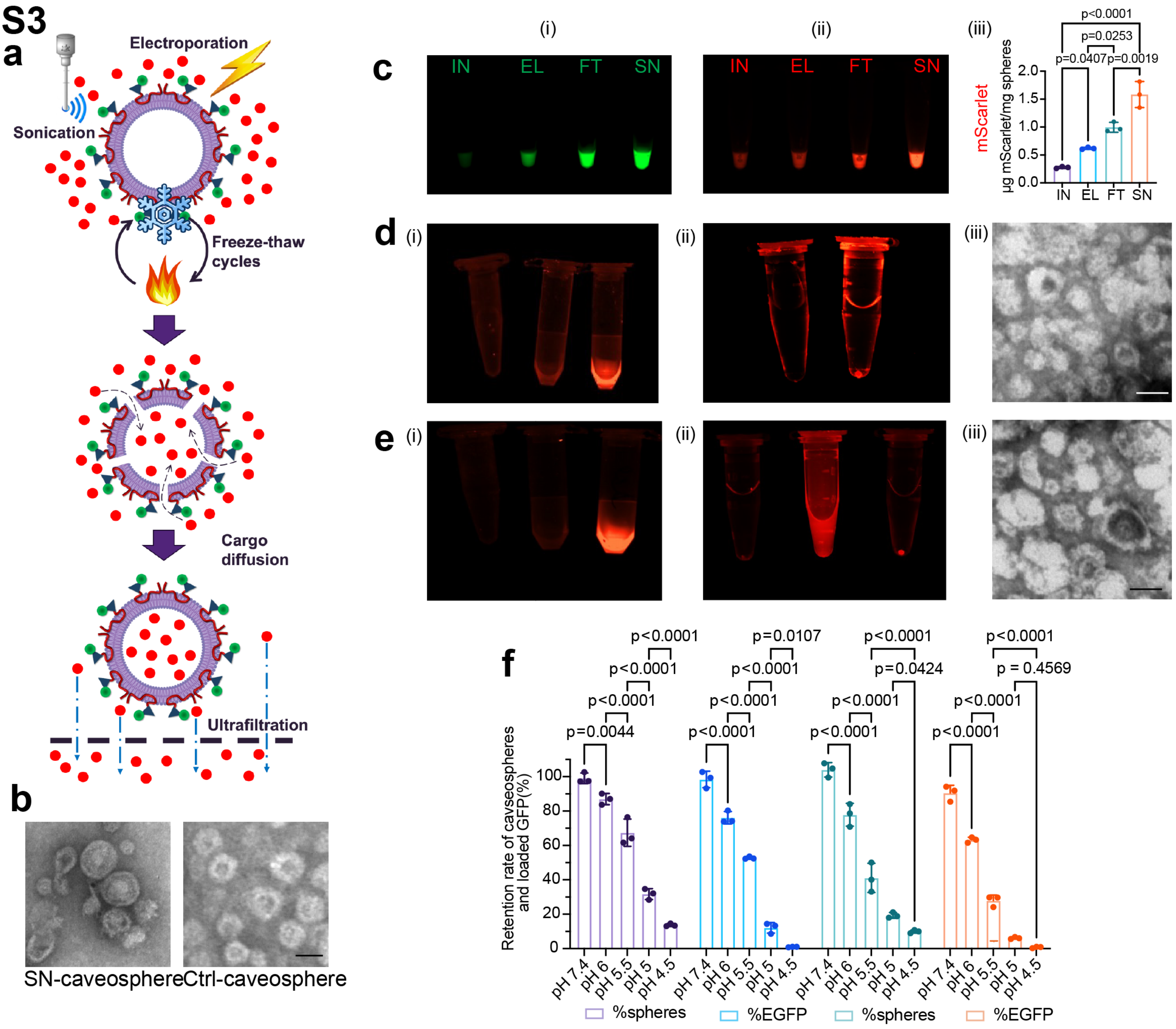

**Supplementary Figure 3: a,** Schematic diagram of three physical methods used for cargo encapsulation into caveospheres (sonication [SN], electroporation [EL] and freeze-thaw cycling [FT]). **b,** TEM comparison of EGFP-caveospheres loaded by sonication and unloaded caveospheres (Ctrl). **c,** Fluorescence emission of EGFP- and mScarlet-loaded caveospheres in microfuge tubes (i) and (ii) followed by quantification of fluorescence emission of mScarlet-loaded caveospheres by plate-reader (iii). IN=incubation of proteins and caveospheres without physical treatment. **d,** Fluorescence emission of caveospheres after loading with mScarlet by electroporation: (i) fluorescent imaging in the red channel of PBS 1x, unloaded caveospheres and electroporated caveospheres after purification (left to right), (ii) fluorescent imaging in the red channel after ultracentrifugation of unloaded caveospheres (left) and electroporated caveospheres (right), (iii) negative stained TEM image of electroporated spheres. **e,** Fluorescence characteristics of caveospheres after loading mScarlet by freeze-thaw method: (i) fluorescent imaging in the red channel of PBS 1x, unloaded caveospheres and freeze-thawed caveospheres after purification (left to right), (ii) fluorescent imaging in the red channel after ultracentrifugation of unloaded caveospheres, freeze-thawed caveospheres + TX-100 and freeze-thawed caveospheres without TX-100 (left to right) (iii) TEM image of freeze-thawed caveospheres. **f,** Stability of SN-caveospheres (containing EGFP) after incubation at different pH values at 37°C, followed by neutralizing and purification, compared to untreated SN-caveospheres (Ctrl) using BCA protein assay and fluorescence emission of EGFP (30 min and 1 hr pH treatment) for quantification of percentage of EGFP and caveosphere remaining after treatment compared to untreated ctrl as 100% (shown as retention rate, %). In all panels: Scale bar, 50 nm. n=3 technical replicates, two-way ANOVA with Tukey's multiple comparisons test. Error bars represent mean±SD.

**S4**

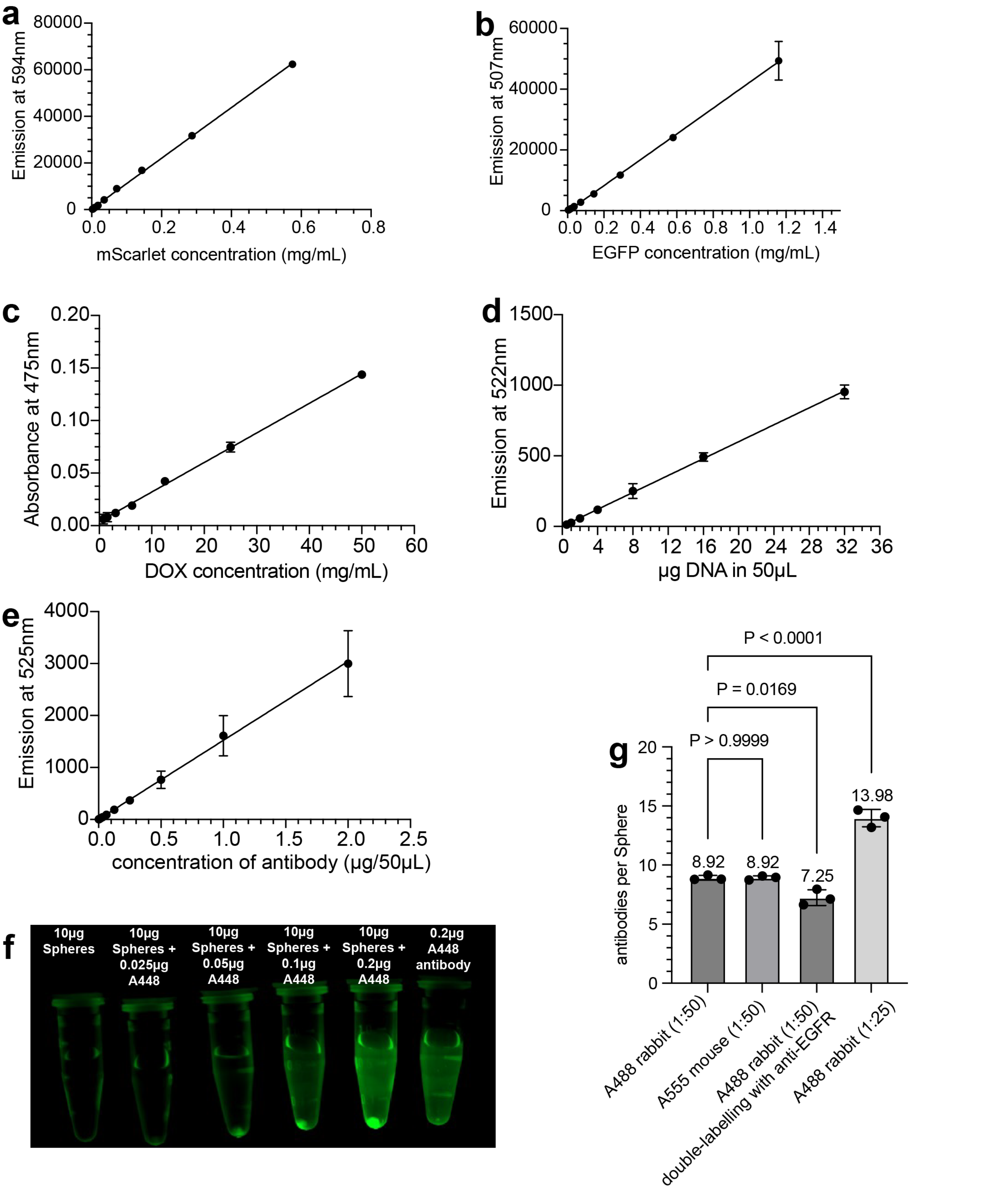

**Supplementary Figure 4: a,** Linear correlation analysis of mScarlet concentration based on fluorescence emission at 594 nm (n=3 technical replicates). **b,** Linear correlation analysis of EGFP concentration based on fluorescence emission at 507 nm (n=3 technical replicates). **c,** Linear correlation analysis between DOX concentration and absorbance at 475 nm (n=5 technical replicates). **d,** Linear correlation analysis of SYBR-labeled DNA concentration and fluorescence emission at 522 nm. (n=5 technical replicates). **e,** Linear correlation analysis between Alexa-488 fluorescent antibody concentration and fluorescence emission at 525 nm (n=3 technical replicates). **f,** Image of fluorescently-labelled spheres in diffirent concentration after ultracentrifugation acquired with ChemiDoc MP imaging system (BIO-RAD) (set at blue epi illumination and 530/28 filter). **g,** Quantification number of A488 antibody, A555 antibody per caveosphere (at 1:50 and 1:25 antibody-to-cavesphere ratio) using the sucrose gradient ultracentrifugation followed by fluorescence emission at 525 nm for A488 and 565 nm for A55 of fractions at 40% and 60% sucrose levels. In all panels: Error bars represent mean±SD.

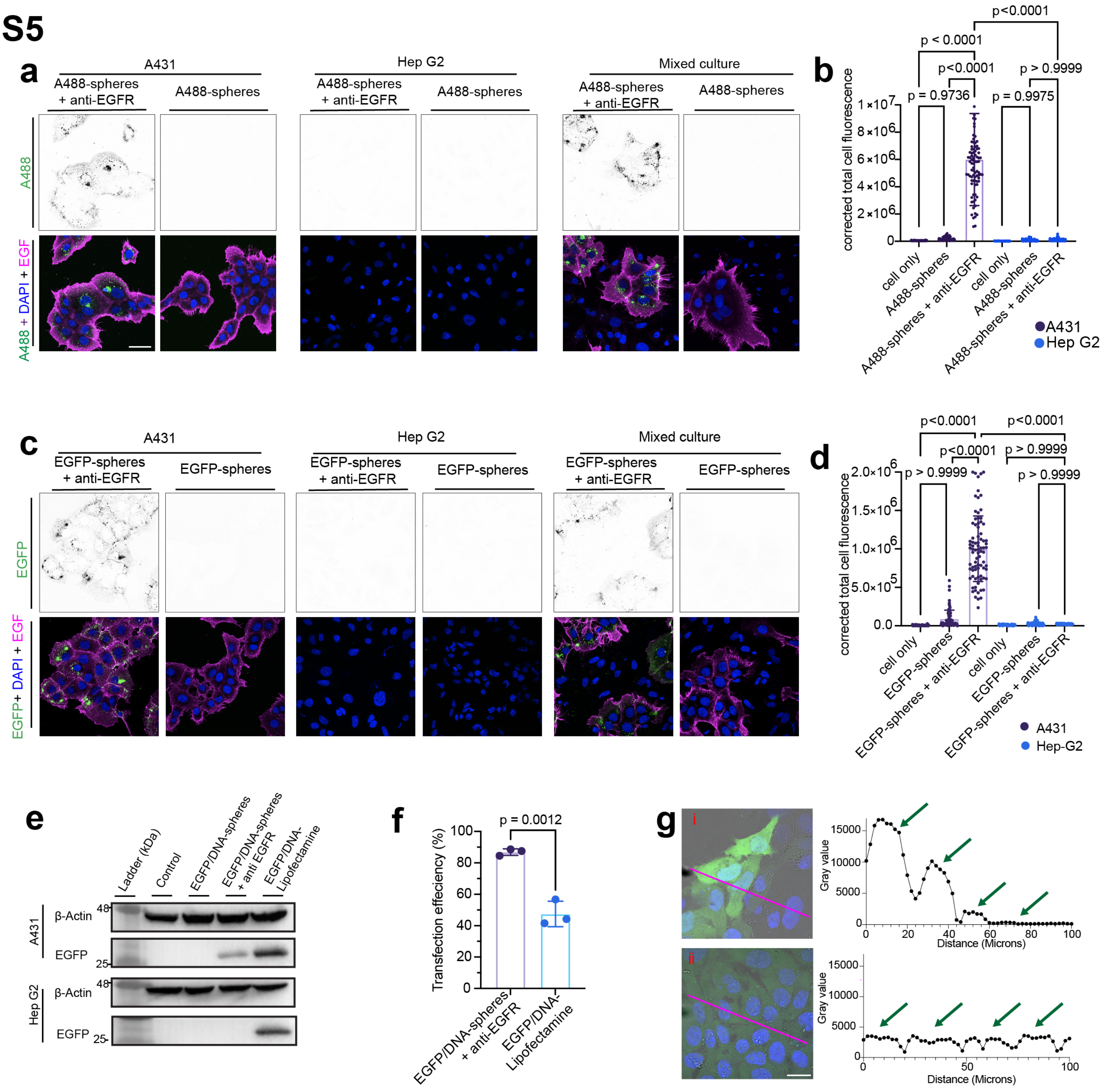

**Supplementary Figure 5: a,** Images of A431, HepG2 and mixed (A431+HepG2) cells after 1 hr uptake of A488-sphere (green) with and without targeting anti-EGFR antibody, stained with Alexa647- conjungated EGF (magenta) and DAPI (blue channel). A488 fluorescence also shown as inverted images. Scale bar, 40 µm. **b,** Quantification (corrected total cell fluorescence) of A488-sphere fluorescence in A431 and HepG2 cells with and without targeting anti-EGFR antibody (n=3 independent experiments, two-way ANOVA with Tukey's multiple comparisons test). **c,** Images of A431, HepG2 and mixed (A431 + HepG2) cells after 1 hr uptake of EGFP-loaded caveospheres (EGFP-spheres; green) with and without targeting anti-EGFR antibody, stained with Alexa647- conjungated EGF (magenta) and DAPI (blue). EGFP fluorescence also shown as inverted images. Scale bar, 40 µm. **d,** Quantification (corrected total cell fluorescence) of EGFP-sphere fluorescence in A431 and HepG2 cells, with and without anti-EGFR antibody (n=3 independent experiments, two-way ANOVA with Tukey's multiple comparison test). **e,** Western analysis of EGFP expression in A431 and Hep G2 cells using EGFP/DNA-spheres +/- anti-EGFR, EGFP/DNA-Lipofectamine, with untreated (Control) cells as a negative control. β-actin is shown as a loading control. **f,** Comparison of transfection efficiency in A431 cells (expressed as percentage of EGFP-positive cells) for EGFP/DNA-spheres + anti-EGFR and EGFP/DNA-Lipofectamine (n=3 independent experiments, student's T-test). **g,** Intensity profile analysis of EGFP expression (gray value) in A431 cells transfected with EGFP/DNA-Lipofectamine (i) or EGFP/DNA-spheres + anti-EGFR (ii). The pixel intensity of EGFP expressed in each cell is indicated by the green arrow. In all panels: Error bars represent mean±SD.

S6

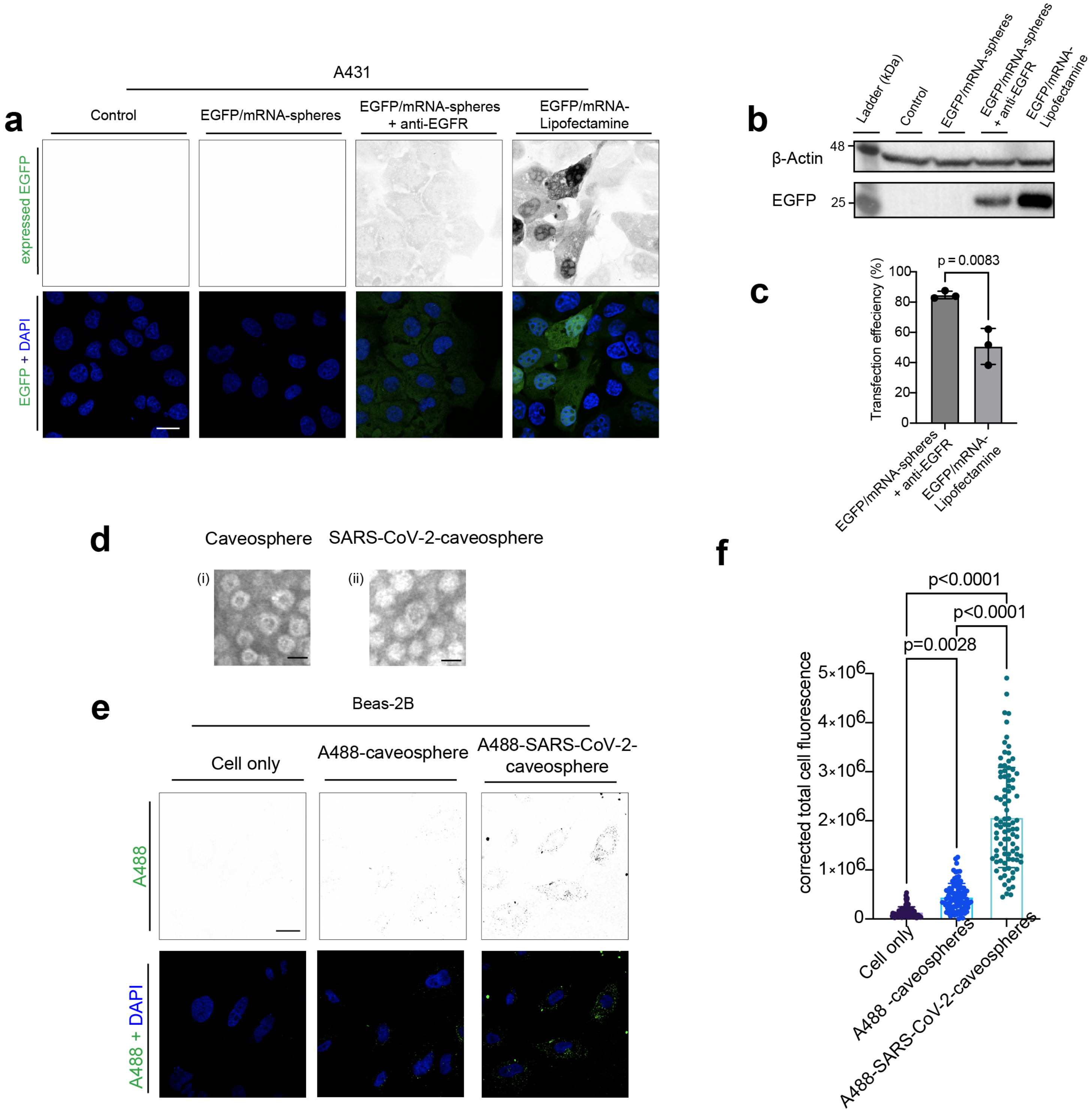

**Supplementary Figure 6:** **a, b** and **c,** A431 cells were transfected with mRNA expressing EGFP loaded caveospheres (EGFP/mRNA-sphere +/- anti-EGFR), in comparison to EGFP/mRNA-Lipofectamine transfection and untreated (Control) cells: **(a)** Imaging of EGFP expression (green) with DAPI staining (blue) in A431 cells. Scale bar, 20 μm. EGFP expression also shown as inverted images. **(b)** Western analysis of EGFP expression in A431 cells, with β-actin as a loading control. **(c)** Comparison of transfection efficiency in A431 cells (expressed as percentage of EGFP-positive cells) for EGFP/mRNA-spheres + anti-EGFR and EGFP/mRNA-Lipofectamine (n=3 independent experiments, total of 90 cells counted, student's T-test). **d,** TEM image of caveospheres (i) and SARS-CoV-2-caveospheres (ii). Scale bars, 50nm. **e,** Images of A488 fluorescence (green) in Beas-2B cells after 3 hrs uptake of A488-caveospheres and A488-SARS-CoV-2-caveospheres, with DAPI staining (blue). A488 fluorescence also shown as inverted images. Scale bar, 20 μm. **f,** Quantification (corrected total cell fluorescence) of A488 fluorescence in Beas-2Bcells after uptake with A488-caveospheres and A488-SARS-CoV-2-caveospheres groups (n=3 independent experiments, one-way ANOVA with Tukey's multiple comparisons test). In all panels: Error bars represent mean±SD.

## S7

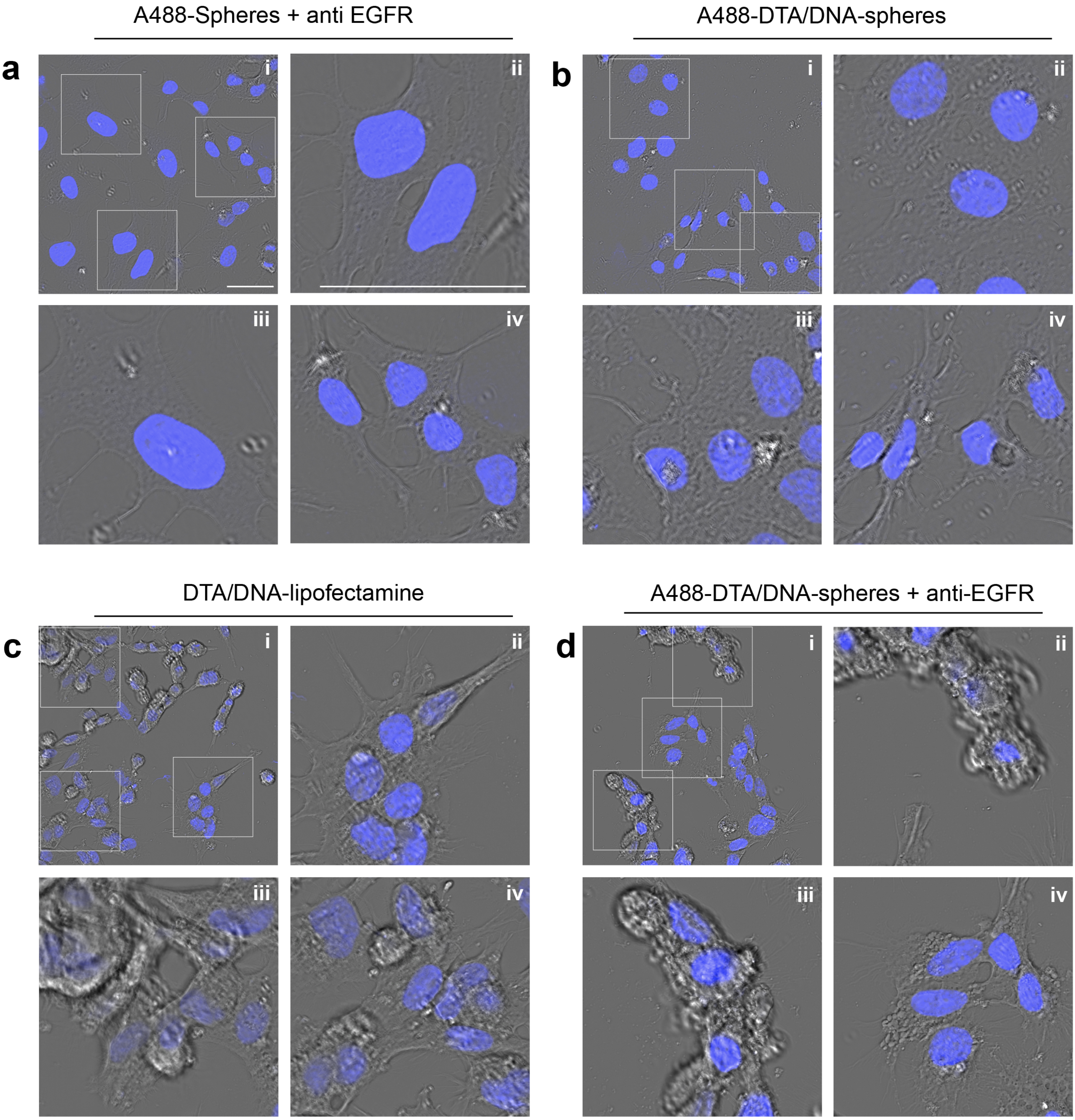

**Supplementary Figure 7:** High-magnification images of A431 and HepG2 cells in mixed culture shown in Figure 2J. **a)** Images of A431 and Hep G2 after 24 hrs incubation with caveospheres labeled with A488 and anti-EGFR antibodies (A488-spheres + anti-EGFR): (i) original image, (ii) and (iii) 3x zoom of the A431 area, (iv) 3x zoom of the HepG2 area. **b)** Images of A431 and Hep G2 after 24 hrs incubation with DNA expressing DTA loaded caveospheres labeled with A488 antibodies without anti-EGFR (A488-DTA/DNA-spheres): (i) original image, (ii) and (iii) 3x zoom of the A431 area, (iv) 3x zoom of the HepG2 area. **c)** Images of A431 and Hep G2 after 24 hrs with DTA transfection with Lipofectamine (DTA/DNA-Lipofectamine): (i) original image, (ii) and (iii) 3x zoom of the A431 area, (iv) 3x zoom of the HepG2 area. **d)** Images of A431 and Hep G2 after 24 hrs incubation with DNA expressing DTA loaded caveospheres labeled with A488 antibodies and anti-EGFR (A488-DTA/DNA-spheres + anti-EGFR): (i) original image, (ii) and (iii) 3x zoom of the A431 area, (iv) 3x zoom of the HepG2 area. Scale bar, 40 µm. Cells were stained with DAPI (blue).

S8

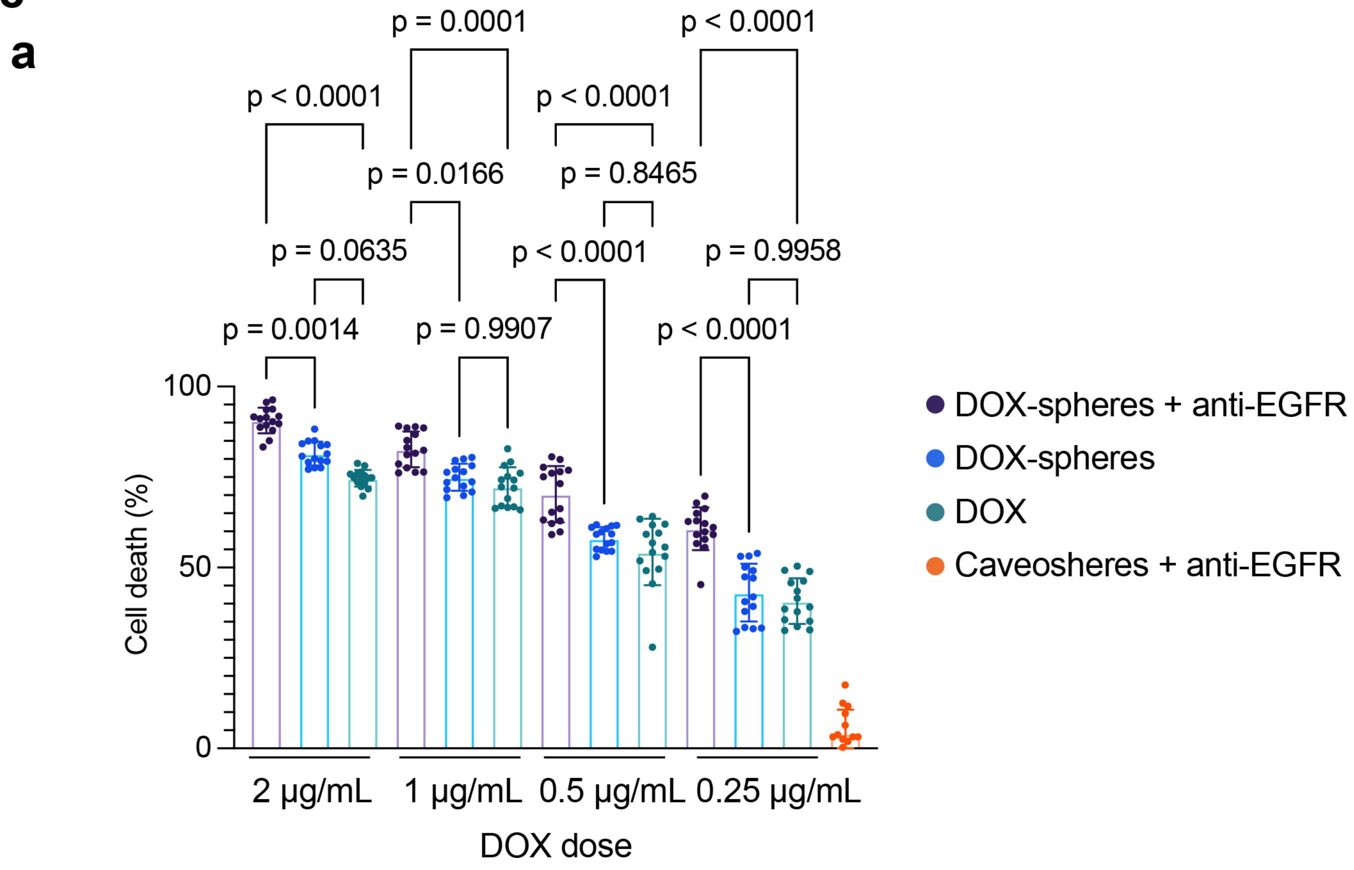

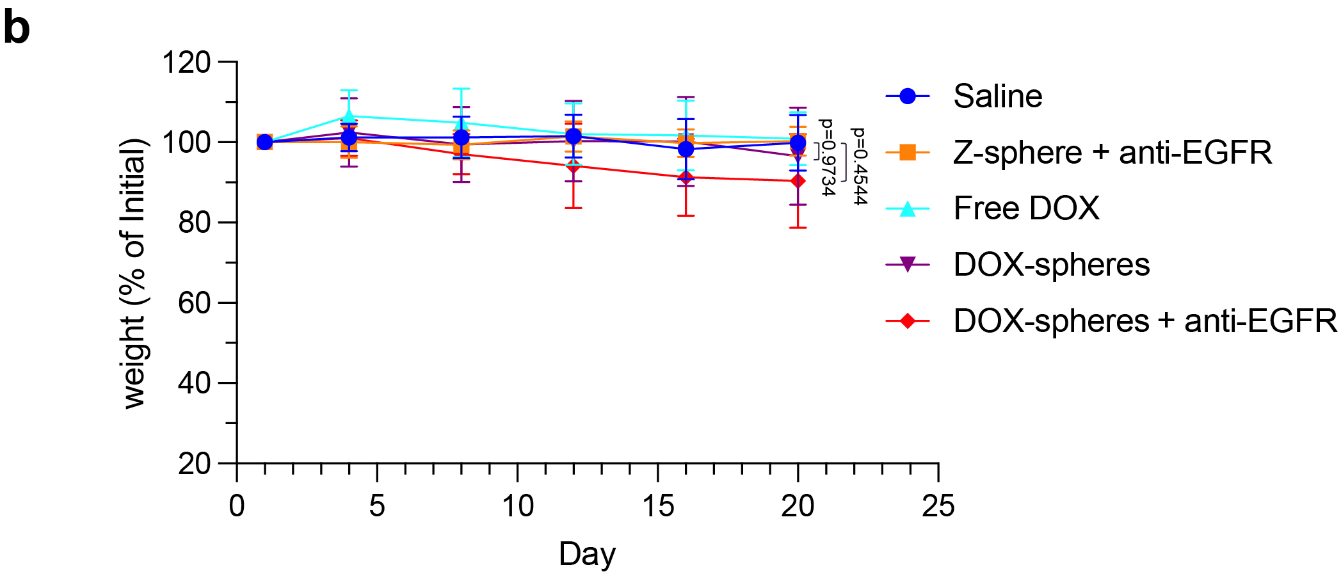

**Supplementary Figure 8: a,** Cell death (%) calculated based on cell viability assay of A431 cells after 48 hrs treatment with DOX-spheres + anti-EGFR, DOX-spheres or free DOX (n=3 independent experiments, two-way ANOVA with Tukey's multiple comparisons test). Untreated cells were set as the 100% survival baseline. **b,** Weight of mice (expressed as a percentage of starting weight at day 1) following a tumor regression study over 20 days (n=5 mice per treatment, one-way ANOVA with Tukey's multiple comparisons test, error bars represent SD). Mice were injected with Free DOX, DOX-spheres or DOX-spheres + anti-EGFR on days 1, 4, 8, 12 and 16 of treatment. Saline and anti-EGFR functionalized caveospheres (-DOX) were used as negative controls. P values between DOX-spheres, DOX-spheres + anti-EGFR and saline provided. In all panels: Error bars represent mean±SD.

S9

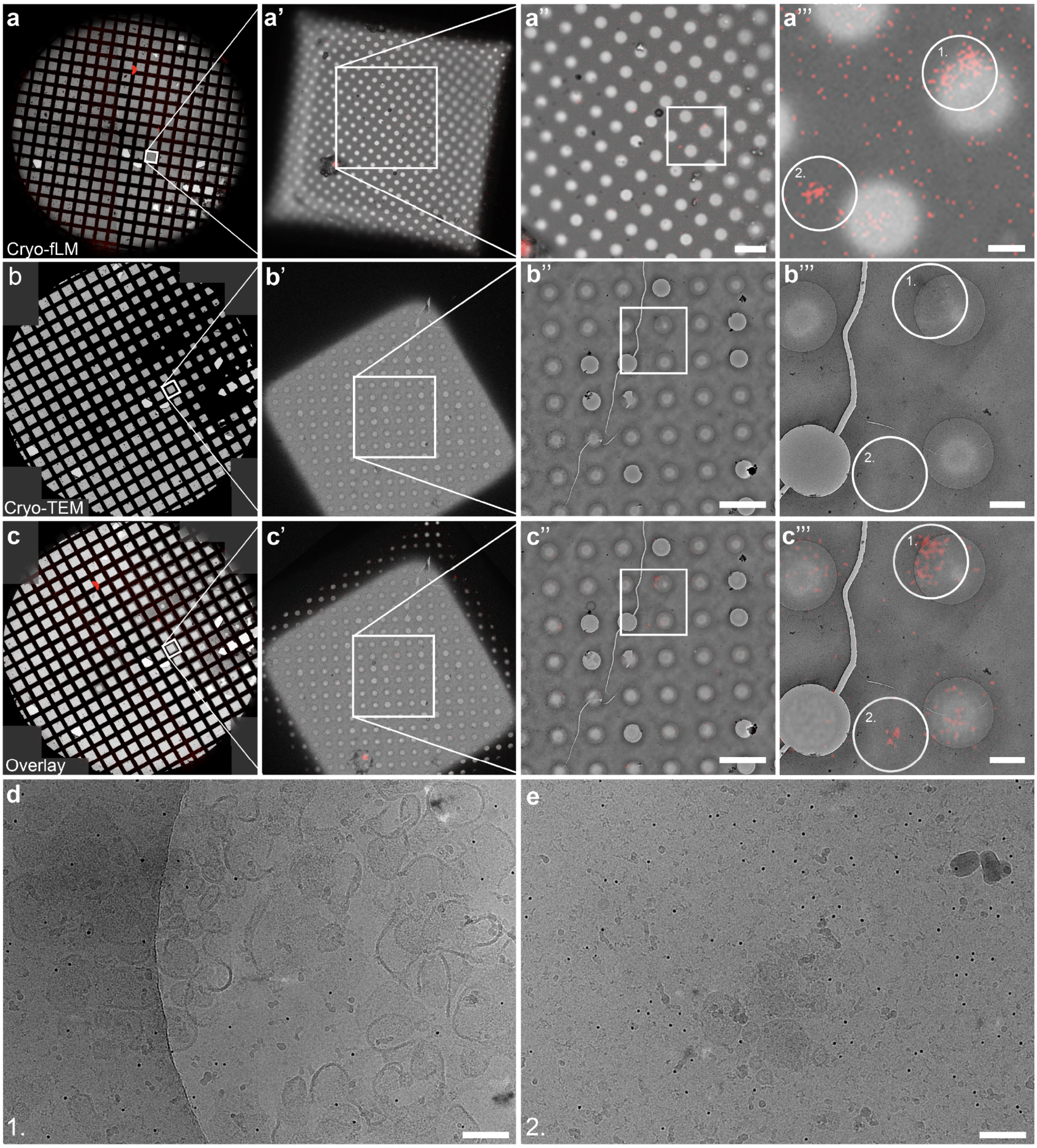

**Supplementary Figure 9:** Validation of the cryo-CLEM approach with additional areas of interest (related to Figure 1h). mScarlet periplasmic expression results in encapsulation of fluorescence within heterologous caveolae. Cryo-fluorescence light microscopy of additional regions of interest shows striking co-localization between mScarlet fluorescence intensity and caveospheres. **a-a'''**, Low magnification brightfield with cryo-confocal detection of the mScarlet signal showing low magnification whole grid **(a)**, to grid square level **(a')**, to region **(a'')**, to hole **(a''')** tracking of mScarlet signal. **b-b'''**, Cryo-EM of detection of the same grid and regions from **(a)**. **c-c''',** Overlay of cryo-fluorescence image from **(a)** with cryo-EM image from **(b)** demonstrating grid fitting **(c)**, to grid square level **(c')**, to region **(c'')**, to hole detection **(c''')** tracking results in the agreement of morphologically definable caveospheres at sites of enriched mScarlet signal. Region 1 and Region 2 are highlighted. **d,** High-magnification of cryo-electron micrograph of region 1 (from **a''', b'''** and **c'''**) shows clustered caveospheres at the area of enriched mScarlet fluorescence. e, High-magnification of cryo-electron micrograph of region 2 (from **a''', b'''** and **c'''**) shows clustered caveospheres at the area of enriched mScarlet fluorescence. Scale bars: 5μm in **a'', b''** and **c''**; 1μm in **a''', b'''** and **c '''**; 100 nm in d and e.

S10

### Supplementary 10: Exact P values and statistics for figure 1

## 1b

| **Unpaired t test** | |
|---|---|
| P value | <0.0001 |
| P value summary | **** |
| Significantly different (P < 0.05)? | Yes |
| One- or two-tailed P value? | Two-tailed |
| t, df | t=23.56, df=4 |
| Difference between means (B - A) ± SEM | 49.67 ± 2.108 |
| 95% confidence interval | 43.81 to 55.52 |

## 1e

| **Tukey's multiple comparisons test** | **Mean Diff.** | **95.00% CI of diff.** | **Summary** | **P Value** | |
|---|---|---|---|---|---|
| In vs. El | -0.4012 | -0.6613 to -0.1411 | ** | 0.005 | |
| In vs. Ft | -0.8042 | -1.064 to -0.5441 | **** | <0.0001 | |
| In vs. Sn | -1.168 | -1.428 to -0.9078 | **** | <0.0001 | |
| El vs. Ft | -0.403 | -0.6631 to -0.1429 | ** | 0.0048 | |
| El vs. Sn | -0.7667 | -1.027 to -0.5066 | **** | <0.0001 | |
| Ft vs. Sn | -0.3637 | -0.6238 to -0.1036 | ** | 0.0089 | |
| **ANOVA summary** | | | | | |
| F | 77.17 | | | | |
| P value | <0.0001 | | | | |
| **ANOVA table** | SS | DF | MS | F (DFn, DFd) | P value |
| Treatment (between columns) | 2.291 | 3 | 0.7636 | F (3, 8) = 77.17 | P<0.0001 |

## 1i

| **Unpaired t test** | |
|---|---|
| P value | <0.0001 |
| P value summary | **** |
| Significantly different (P < 0.05)? | Yes |
| One- or two-tailed P value? | Two-tailed |
| t, df | t=20.27, df=8 |
| Difference between means (B - A) ± SEM | 30.60 ± 1.510 |
| 95% confidence interval | 27.12 to 34.08 |

## 1o

| **Unpaired t test** | |
|---|---|
| P value | <0.0001 |
| P value summary | **** |
| Significantly different (P < 0.05)? | Yes |
| One- or two-tailed P value? | Two-tailed |
| t, df | t=19.23, df=4 |
| Difference between means (B - A) ± SEM | 1.994 ± 0.1037 |
| 95% confidence interval | 1.706 to 2.282 |

## 1t

| **Unpaired t test** | |
|---|---|
| P value | 0.9702 |
| P value summary | ns |
| Significantly different (P < 0.05)? | No |
| One- or two-tailed P value? | Two-tailed |
| t, df | t=0.03970, df=4 |
| Difference between means (B - A) ± SEM | -0.01392 ± 0.3505 |
| 95% confidence interval | -0.9871 to 0.9592 |

### S11 Supplementary 11: Exact P values and statistics for figure 2

## 2b

| Tukey's multiple comparisons test | Mean Diff. | 95.00% CI of diff. | Summary | P Value | |
|---|---|---|---|---|---|
| A431 | | | | | |
| Control vs. EGFP/DNA-spheres | -100879 | -1787196 to 1585439 | ns | >0.9999 | |
| Control vs. EGFP/DNA-spheres + anti EGFR | -2479453 | -4206636 to -752270 | *** | 0.0004 | |
| Control vs. Lipofectamine | -7225328 | -8975787 to -5474869 | **** | <0.0001 | |
| EGFP/DNA-spheres vs. EGFP/DNA-spheres + anti EGFR | -2378574 | -4105757 to -651392 | *** | 0.0008 | |
| EGFP/DNA-spheres vs. Lipofectamine | -7124449 | -8874908 to -5373990 | **** | <0.0001 | |
| HepG2 | | | | | |
| Control vs. EGFP/DNA-spheres | 10785 | -1746762 to 1768331 | ns | >0.9999 | |
| Control vs. EGFP/DNA-spheres + anti EGFR | -301247 | -2043700 to 1441206 | ns | 0.9995 | |
| Control vs. Lipofectamine | -6112236 | -7815886 to -4408586 | **** | <0.0001 | |
| EGFP/DNA-spheres vs. EGFP/DNA-spheres + anti EGFR | -312032 | -2123545 to 1499482 | ns | 0.9995 | |
| EGFP/DNA-spheres vs. Lipofectamine | -6123021 | -7893159 to -4352883 | **** | <0.0001 | |
| A431 vs. HepG2 | | | | | |
| EGFP/DNA-spheres + anti EGFR vs. EGFP/DNA-spheres + anti EGFR | 2146936 | 372642 to 3921231 | ** | 0.0062 | |
| Lipofectamine vs. Lipofectamine | 1081822 | -640371 to 2804015 | ns | 0.544 | |
| **Source of Variation** | **% of total variation** | **P value** | **P value summary** | **Significant?** | |
| Row Factor | 20.58 | 0.0082 | ** | Yes | |
| Column Factor | 30.28 | <0.0001 | **** | Yes | |
| **ANOVA table** | **SS (Type III)** | **DF** | **MS** | **F (DFn, DFd)** | **P value** |
| Row Factor | 3.423E+15 | 188 | 1.821E+13 | F (188, 596) = 1.317 | P=0.0082 |
| Column Factor | 5.038E+15 | 7 | 7.197E+14 | F (7, 596) = 52.03 | P<0.0001 |
| Residual | 8.243E+15 | 596 | 1.3831E+13 | | |

## 2f

| Tukey's multiple comparisons test | Mean Diff. | 95.00% CI of diff. | Summary | P Value | |
|---|---|---|---|---|---|
| Control vs. EGFP/DNA-spheres | -38713 | -194752 to 117327 | ns | 0.8284 | |
| Control vs. EGFP/DNA-SARS-CoV-2-caveosphere | -979759 | -1134130 to -825389 | **** | <0.0001 | |
| EGFP/DNA-spheres vs. EGFP/DNA-SARS-CoV-2-caveosphere | -941047 | -1095417 to -786676 | **** | <0.0001 | |
| **ANOVA summary** | | | | | |
| F | 144.5 | | | | |
| P value | <0.0001 | | | | |
| P value summary | **** | | | | |
| Significant diff. among means (P < 0.05)? | Yes | | | | |
| **ANOVA table** | **SS** | **DF** | **MS** | **F (DFn, DFd)** | **P value** |
| Treatment (between columns) | 5.70257E+13 | 2 | 2.8513E+13 | F (2, 271) = 144.5 | P<0.0001 |
| Residual (within columns) | 5.34632E+13 | 271 | 1.9728E+11 | | |

## 2h

| Tukey's multiple comparisons test | Mean Diff. | 95.00% CI of diff. | Summary | Adjusted P Value | |
|---|---|---|---|---|---|
| control vs. minus anti-CD3e | -34803 | -128363 to 58756 | ns | 0.7734 | |
| control vs. plus anti-CD3e | -602179 | -690462 to -513895 | **** | <0.0001 | |
| control vs. lipofectamine | -173081 | -262890 to -83271 | **** | <0.0001 | |
| minus anti-CD3e vs. plus anti-CD3e | -567375 | -649113 to -485638 | **** | <0.0001 | |
| minus anti-CD3e vs. lipofectamine | -138278 | -221661 to -54894 | *** | 0.0001 | |
| plus anti-CD3e vs. lipofectamine | 429098 | 351681 to 506514 | **** | <0.0001 | |
| **ANOVA summary** | | | | | |
| F | 154.7 | | | | |
| P value | <0.0001 | | | | |
| **ANOVA table** | SS | DF | MS | F (DFn, DFd) | P value |
| Treatment (between columns) | 4.3839E+13 | 3 | 1.4613E+13 | F (3, 706) = 154.7 | P<0.0001 |

## 2i

| Unpaired t test | |
|---|---|
| P value | <0.0001 |
| P value summary | **** |
| Significantly different (P < 0.05)? | Yes |
| One- or two-tailed P value? | Two-tailed |
| t, df | t=165.5, df=4 |
| Difference between means (B - A) ± SEM | -77.05 ± 0.4655 |
| 95% confidence interval | -78.34 to -75.76 |

### S12 Supplementary 12: Exact P values and statistics for figure 2

## 2k

| Tukey's multiple comparisons test | Mean Diff. | 95.00% CI of diff. | Summary | P Value | |
|---|---|---|---|---|---|
| DTA/DNA-Sphere + anti-EGFR A431 vs. DTA/DNA-Sphere A431 | 71.46 | 61.05 to 81.87 | **** | <0.0001 | |
| DTA/DNA-Sphere + anti-EGFR A431 vs. DTA/DNA-lipofectamine 431 | 29.5 | 19.09 to 39.91 | **** | <0.0001 | |
| DTA/DNA-Sphere + anti-EGFR A431 vs. EGFP/DNA-Sphere + anti-EGFR A431 | 66.65 | 56.24 to 77.06 | **** | <0.0001 | |
| DTA/DNA-Sphere + anti-EGFR A431 vs. DTA/DNA-Sphere + anti-EGFR HepG2 | 73.26 | 62.85 to 83.67 | **** | <0.0001 | |
| DTA/DNA-lipofectamine A431 vs. EGFP/DNA-lipofectamine A431 | 33.62 | 23.21 to 44.03 | **** | <0.0001 | |
| DTA/DNA-lipofectamine A431 vs. DTA/DNA-lipofectamine HepG2 | 33.37 | 22.96 to 43.78 | ns | 0.9888 | |
| EGFP/DNA-Sphere + anti-EGFR A431 vs. EGFP/DNA-Sphere A431 | 8.264 | -2.145 to 18.67 | ns | 0.2699 | |
| EGFP/DNA-Sphere + anti-EGFR A431 vs. EGFP/DNA-lipofectamine A431 | -3.533 | -13.94 to 6.877 | ns | 0.993 | |
| DTA/DNA-Sphere + anti-EGFR HepG2 vs. DTA/DNA-Sphere HepG2 | 3.799 | -6.611 to 14.21 | ns | 0.9874 | |
| DTA/DNA-Sphere + anti-EGFR HepG2 vs. DTA/DNA-lipofectamine HepG2 | -40.01 | -50.42 to -29.60 | **** | <0.0001 | |
| DTA/DNA-Sphere + anti-EGFR HepG2 vs. GFP/DNA-Sphere + anti-EGFR HepG2 | 1.789 | -8.621 to 12.20 | ns | >0.9999 | |
| DTA/DNA-lipofectamine HepG2 vs. GFP/DNA-lipofectamine HepG2 | 29.63 | 19.22 to 40.04 | **** | <0.0001 | |
| GFP/DNA-Sphere + anti-EGFR HepG2 vs. GFP/DNA-Sphere HepG2 | -0.6557 | -11.07 to 9.754 | ns | >0.9999 | |
| GFP/DNA-Sphere + anti-EGFR HepG2 vs. GFP/DNA-lipofectamine HepG2 | -12.17 | -22.58 to -1.762 | ** | 0.0082 | |
| GFP/DNA-Sphere HepG2 vs. GFP/DNA-lipofectamine HepG2 | -11.52 | -21.93 to -1.106 | * | 0.0167 | |
| **Source of Variation** | **% of total variation** | **P value** | **P value summary** | **Significant?** | |
| Row Factor | 1.699 | 0.0387 | * | Yes | |
| Column Factor | 88.09 | <0.0001 | **** | Yes | |
| **ANOVA table** | **SS (Type III)** | **DF** | **MS** | **F (DFn, DFd)** | **P value** |
| Row Factor | 1891 | 14 | 135 | F (14, 154) = 1.830 | P=0.0387 |
| Column Factor | 98002 | 11 | 8909 | F (11, 154) = 120.8 | P<0.0001 |
| Residual | 11362 | 154 | 73.78 | | |

### S13 Supplementary 13: Exact P values and statistics for figure 3

## 3c

| Šídák's multiple comparisons test | Mean Diff. | 95.00% CI of diff. | Summary | P Value | |
|---|---|---|---|---|---|
| A680-sphere vs. A680-sphere + anti-EGFR | | | | | |
| Liver | -30613400 | -277866934 to 216640134 | ns | >0.9999 | |
| Tumour | -393061400 | -640314934 to -145807866 | *** | 0.0002 | |
| Blood | -28489400 | -275742934 to 218764134 | ns | >0.9999 | |
| GI | 16772200 | -230481334 to 264025734 | ns | >0.9999 | |
| Lungs | -4660200 | -251913734 to 242593334 | ns | >0.9999 | |
| Kidneys | -34215400 | -281468934 to 213038134 | ns | >0.9999 | |
| Spleen | -4712200 | -251965734 to 242541334 | ns | >0.9999 | |
| Heart | -59901400 | -307154934 to 187352134 | ns | 0.9959 | |
| **Source of Variation** | **% of total variation** | **P value** | **P value summary** | **Significant?** | |
| Interaction | 6.039 | 0.0355 | * | Yes | |
| Row Factor | 68.44 | <0.0001 | **** | Yes | |
| Column Factor | 1.753 | 0.0335 | * | Yes | |
| **ANOVA table** | **SS** | **DF** | **MS** | **F (DFn, DFd)** | **P value** |
| Interaction | 3.126E+17 | 7 | 4.465E+16 | F (7, 64) = 2.323 | P=0.0355 |
| Row Factor | 3.542E+18 | 7 | 5.06E+17 | F (7, 64) = 26.32 | P<0.0001 |
| Column Factor | 9.075E+16 | 1 | 9.075E+16 | F (1, 64) = 4.721 | P=0.0335 |
| Residual | 1.23E+18 | 64 | 1.922E+16 | | |

## 3f

| Unpaired t test | |
|---|---|
| P value | 0.001 |
| P value summary | *** |
| Significantly different (P < 0.05)? | Yes |
| One- or two-tailed P value? | Two-tailed |
| t, df | t=5.043, df=8 |
| Difference between means (C - B) ± SEM | 7820 ± 1551 |
| 95% confidence interval | 4245 to 11396 |

### S14 Supplementary 14: Exact P values and statistics for figure 4

## 4b

| Tukey's multiple comparisons test | Mean Diff. | 95.00% CI of diff. | Summary | P Value | |
|---|---|---|---|---|---|
| DOX in nucleus (red Chanel) | | | | | |
| 30 mins: | | | | | |
| DOX-spheres + anti-EGFR vs. DOX-spheres | 1827547 | 1344174 to 2310921 | **** | <0.0001 | |
| DOX-spheres + anti-EGFR vs. DOX | 1822293 | 1319224 to 2325362 | **** | <0.0001 | |
| DOX-spheres vs. DOX | 2194695 | 1690044 to 2699346 | ns | >0.9999 | |
| 1 hr: | | | | | |
| DOX-spheres + anti-EGFR vs. DOX-spheres | 2561000 | 2073464 to 3048535 | **** | <0.0001 | |
| DOX-spheres + anti-EGFR vs. DOX | 2600822 | 2100699 to 3100945 | **** | <0.0001 | |
| DOX-spheres vs. DOX | 39823 | -454866 to 534511 | ns | >0.9999 | |
| 2hrs: | | | | | |
| DOX-spheres + anti-EGFR vs. DOX-spheres | 3972881 | 3477978 to 4467784 | **** | <0.0001 | |
| DOX-spheres + anti-EGFR vs. DOX-spheres | 4227665 | 3737235 to 4718095 | **** | <0.0001 | |
| DOX-spheres vs. DOX | 1241517 | 739544 to 1743490 | ns | 0.9509 | |
| 488 antibody in cytoplasm (green channel) | | | | | |
| 30 mins: | | | | | |
| DOX-spheres + anti-EGFR vs. DOX-spheres | 1706306 | 1223090 to 2189521 | **** | <0.0001 | |
| DOX-spheres + anti-EGFR vs. DOX | 1875534 | 1369108 to 2381961 | **** | <0.0001 | |
| DOX-spheres vs. DOX | 169229 | -328790 to 667247 | ns | 0.9994 | |
| 1 hr: | | | | | |
| DOX-spheres + anti-EGFR vs. DOX-spheres | 2732048 | 2244484 to 3219613 | **** | <0.0001 | |
| DOX-spheres + anti-EGFR vs. DOX | 3243231 | 2750310 to 3736152 | **** | <0.0001 | |
| DOX-spheres vs. DOX | 511183 | 23801 to 998564 | * | 0.0283 | |
| 2hrs: | | | | | |
| DOX-spheres + anti-EGFR vs. DOX-spheres | 3535157 | 3040254 to 4030060 | **** | <0.0001 | |
| DOX-spheres + anti-EGFR vs. DOX-spheres | 4419400 | 3930604 to 4908195 | **** | <0.0001 | |
| DOX-spheres vs. DOX | 884243 | 382270 to 1386216 | **** | <0.0001 | |
| **Source of Variation** | **% of total variation** | **P value** | **P value summary** | **Significant?** | |
| Row Factor | 1.842 | 0.5508 | ns | No | |
| Column Factor | 71.27 | <0.0001 | **** | Yes | |
| **ANOVA table** | **SS (Type III)** | **DF** | **MS** | **F (DFn, DFd)** | **P value** |
| Row Factor | 1.089E+14 | 117 | 9.30914E+11 | F (117, 1604) = 0.9776 | P=0.5508 |
| Column Factor | 4.214E+15 | 17 | 2.479E+14 | F (17, 1604) = 260.3 | P<0.0001 |

## 4d

| Tukey's multiple comparisons test | Mean Diff. | diff. | Summary | P Value | |
|---|---|---|---|---|---|
| Saline vs. Caveosphere + anti-EGFR | 16.01 | -35.57 to 67.58 | ns | 0.8824 | |
| Saline vs. Free DOX | 105.3 | 53.76 to 156.9 | **** | <0.0001 | |
| Saline vs. DOX-sphere | 128.3 | 76.72 to 179.9 | **** | <0.0001 | |
| Saline vs. DOX-sphere + anti-EGFR | 195.3 | 143.7 to 246.9 | **** | <0.0001 | |
| Caveosphere + anti-EGFR vs. Free DOX | 89.33 | 37.76 to 140.9 | *** | 0.0004 | |
| Caveosphere + anti-EGFR vs. DOX-sphere | 112.3 | 60.71 to 163.9 | **** | <0.0001 | |
| Caveosphere + anti-EGFR vs. DOX-sphere + anti-EGFR | 179.3 | 127.7 to 230.8 | **** | <0.0001 | |
| Free DOX vs. DOX-sphere | 22.96 | -28.61 to 74.53 | ns | 0.6753 | |
| Free DOX vs. DOX-sphere + anti-EGFR | 89.94 | 38.37 to 141.5 | *** | 0.0004 | |
| DOX-sphere vs. DOX-sphere + anti-EGFR | 66.98 | 15.41 to 118.6 | ** | 0.0073 | |
| **ANOVA summary** | | | | | |
| F | 44.36 | | | | |
| P value | <0.0001 | | | | |
| P value summary | **** | | | | |
| Significant diff. among means (P < 0.05)? | Yes | | | | |
| **ANOVA table** | **SS** | **DF** | **MS** | **F (DFn, DFd)** | **P value** |
| Treatment (between columns) | 131773 | 4 | 32943 | F (4, 20) = 44.36 | P<0.0001 |
| Residual (within columns) | 14852 | 20 | 742.6 | | |
| Total | 146625 | 24 | | | |

### S15 Supplementary 15: Exact P values and statistics for figure 4

## 4f

| Tukey's multiple comparisons test | Mean Diff. | 95.00% CI of diff. | Summary | P Value | |
|---|---|---|---|---|---|
| Caveosphere + anti-EGFR vs. Free DOX | -62.56 | -75.20 to -49.92 | **** | <0.0001 | |
| Caveosphere + anti-EGFR vs. DOX-sphere | -63.89 | -76.52 to -51.25 | **** | <0.0001 | |
| Caveosphere + anti-EGFR vs. DOX-sphere + anti-EGFR | -89.5 | -102.1 to -76.86 | **** | <0.0001 | |
| Free DOX vs. DOX-sphere | -1.326 | -13.97 to 11.31 | ns | 0.9902 | |
| Free DOX vs. DOX-sphere + anti-EGFR | -26.94 | -39.58 to -14.30 | **** | <0.0001 | |
| DOX-sphere vs. DOX-sphere + anti-EGFR | -25.61 | -38.25 to -12.97 | *** | 0.0001 | |
| **ANOVA summary** | | | | | |
| F | 148.5 | | | | |
| P value | <0.0001 | | | | |
| P value summary | **** | | | | |
| Significant diff. among means (P < 0.05)? | Yes | | | | |
| **ANOVA table** | **SS** | **DF** | **MS** | **F (DFn, DFd)** | **P value** |
| Treatment (between columns) | 21736 | 3 | 7245 | F (3, 16) = 148.5 | P<0.0001 |
| Residual (within columns) | 780.7 | 16 | 48.8 | | |
| Total | 22517 | 19 | | | |

## 4h

| Tukey's multiple comparisons test | Mean Diff. | 95.00% CI of diff. | Summary | P Value | |
|---|---|---|---|---|---|
| PBS 1x vs. Free DOX | 365.2 | 198.0 to 532.4 | *** | 0.0002 | |
| PBS 1x vs. DOX-Sphere + EGFR | 53.04 | -114.2 to 220.2 | ns | 0.6827 | |
| Free DOX vs. DOX-Sphere + EGFR | -312.1 | -479.4 to -144.9 | *** | 0.0009 | |
| **ANOVA summary** | | | | | |
| F | 19.82 | | | | |
| P value | 0.0002 | | | | |
| P value summary | *** | | | | |
| Significant diff. among means (P < 0.05)? | Yes | | | | |
| **ANOVA table** | **SS** | **DF** | **MS** | **F (DFn, DFd)** | **P value** |
| Treatment (between columns) | 389352 | 2 | 194676 | F (2, 12) = 19.82 | P=0.0002 |
| Residual (within columns) | 117839 | 12 | 9820 | | |
| Total | 507191 | 14 | | | |

S16

#### Supplementary 15: Exact P values and statistics for figure 5

**5e**

| **Unpaired t test** | |
|---|---|
| P value | 0.0005 |
| P value summary | *** |
| Significantly different ($P < 0.05$)? | Yes |
| One- or two-tailed P value? | Two-tailed |
| t, df | t=10.19, df=4 |
| 95% confidence interval | 4048 to 7079 |
